# *Helicobacter pylori* provokes STING immunosurveillance via trans-kingdom conjugation

**DOI:** 10.1101/2022.06.29.498044

**Authors:** Prashant P. Damke, Cecily R. Wood, Carrie L. Shaffer

## Abstract

Recognition of foreign nucleic acids is an evolutionarily conserved mechanism by which the host detects microbial threats. Whereas some intracellular bacterial pathogens trigger DNA surveillance pathways following phagosomal membrane perturbation, mechanisms by which extracellular bacteria activate cytosolic nucleic acid reconnaissance systems remain unresolved. Here, we demonstrate that *Helicobacter pylori* exploits *cag* type IV secretion system (*cag* T4SS) activity to provoke STING signaling in gastric epithelial cells. We provide direct evidence that chromosomal fragments delivered to the host cell cytoplasm via trans-kingdom conjugation bind and activate the key DNA sensor cGMP-AMP synthase. To enable paracrine-like signal amplification, translocated *H. pylori* DNA is sorted into exosomes that stimulate DNA-sensing pathways in uninfected bystander cells. We show that DNA cargo is loaded into the *cag* T4SS apparatus in the absence of host cell contact to establish a ‘ready-to-fire’ nanomachine and provide evidence that *cag* T4SS-dependent DNA translocation is mechanistically coupled to chromosomal replication and replichore decatenation. Collectively, these studies suggest that *H. pylori* evolved mechanisms to stimulate nucleic acid surveillance pathways that regulate both pro- and anti-inflammatory programs to facilitate chronic persistence in the gastric niche.

## INTRODUCTION

Innate recognition of invariant pathogen-associated molecular signatures by cellular pattern recognition receptors (PRRs) is the first line of defense against microbial adversaries. Within the gastrointestinal tract, epithelial cells express proximal endosomal and cytosolic nucleic acid-sensing PRRs that rapidly respond to aberrant microbial DNA and RNA to trigger innate defense mechanisms and coordinate adaptative immunity. Localization of foreign DNA to the host cell cytosol activates multiple nucleic acid reconnaissance systems including the key DNA sensor nucleotidyltransferase cyclic GMP-AMP synthase (cGAS) (Cai et al., 2014; Diner et al., 2013; Sun et al., 2013; Wu et al., 2013). Upon binding DNA originating from either extrinsic or intrinsic sources, cGAS catalyzes the formation of the non-canonical cyclic di-nucleotide 2’3’-cGAMP using ATP and GTP as substrates (Diner et al., 2013; Gao et al., 2013a; Gao et al., 2013b; Sun et al., 2013; Zhang et al., 2013). In turn, 2’3’-cGAMP stimulates the endoplasmic reticulum receptor STING (<u>st</u>imulator of <u>in</u>terferon <u>g</u>enes) to elicit interferon (IFN) signaling and the production of multifarious inflammatory cytokines (Ishii et al., 2006; Ishikawa and Barber, 2008; Ishikawa et al., 2009; Stetson and Medzhitov, 2006; Sun et al., 2009; Zhong et al., 2008). Dysregulation of mucosal STING signaling can disrupt gut homeostasis and generate pro-tumorigenic inflammatory microenvironments (Ke et al., 2022); however, the outcomes of STING-dependent immune surveillance within the context of gastric inflammation and infection-associated carcinogenesis remain unresolved.

Of the known infection-associated cancers, the most significant carcinogenic microbe is the gastric bacterium *Helicobacter pylori*, which chronically colonizes the stomach of over half of the global population and directly contributes to the development of more than one million new cases of cancer per year (Sung et al., 2021). *H. pylori* harboring the cancer-associated *cag* type IV secretion system (*cag* T4SS) significantly augment disease risk via translocation of pro-inflammatory molecular cargo into gastric epithelial cells. In addition to facilitating the delivery of the bacterial oncoprotein CagA, the *cag* T4SS translocates a diverse repertoire of immunostimulatory lipid, nucleic acid, and polysaccharide substrates directly into the gastric epithelium (Amieva and Peek, 2016; Cover et al., 2020). Recent work reported that *H. pylori cag* T4SS activity activates the endosomal DNA-sensing PRR Toll-like Receptor 9 (TLR9), leading to immune suppression conferring tolerance (Varga et al., 2016a; Varga et al., 2016b) as well as other potential inflammation-independent carcinogenic phenotypes (Castano-Rodriguez et al., 2014). In addition to TLR9, multiple cellular nucleic acid sensors, including cGAS (Ablasser et al., 2013; Cai et al., 2014; Diner et al., 2013; Gao et al., 2015; Nandakumar et al., 2019; Storek et al., 2015; Watson et al., 2015; Zhang et al., 2014), RIG-I (Chow et al., 2015; Dixit and Kagan, 2013; Onomoto et al., 2021; Rad et al., 2009), MDA5 (Dixit and Kagan, 2013; Wu et al., 2020), AIM2 (Rathinam et al., 2010), ZBP1/DAI (Kuriakose et al., 2016), IFI-16 (Almine et al., 2017; Unterholzner et al., 2010), and RNA pol III (Ablasser et al., 2009; Chiu et al., 2009) are expressed in the human gastric epithelium and associated mucosal dendritic cells (Rad et al., 2009), raising the hypothesis that *H. pylori* stimulates additional innate nucleic acid surveillance pathways.

While intracellular bacterial pathogens elicit cGAS-STING signaling following phagosomal membrane destabilization or rupture achieved in a type III, IV, VI, or VII secretion system-dependent manner (Ku et al., 2020; Marinho et al., 2017; Nandakumar et al., 2019; Storek et al., 2015; Watson et al., 2015; Zhang et al., 2014), the mechanisms by which extracellular bacteria stimulate DNA reconnaissance systems remain unclear. STING signaling has been implicated in gastric carcinogenesis and *H. pylori* has been shown to activate STING *in vivo* (Song et al., 2017), but whether *H. pylori*-driven STING activation requires cytosolic nucleic acid immunosurvellience is unknown. Here, we demonstrate that *H. pylori*-induced STING signaling is a direct consequence of *cag* T4SS-dependent DNA translocation. We show that *H. pylori* chromosomal fragments delivered to the gastric epithelium via trans-kingdom conjugation directly bind and activate cGAS to stimulate STING signaling. We discovered that upon translocation into primary gastric epithelial cells, fragmented *H. pylori* DNA is sorted into exosomes that are released to amplify foreign nucleic acid immune surveillance in uninfected bystander cells. We provide direct evidence that *cag* T4SS-mediated DNA translocation is mechanistically coupled to chromosomal replication and demonstrate that eukaryotic-optimized constructs greater than 1.5 kb are delivered to the gastric epithelium via *cag* T4SS mechanisms. Our results highlight how *H. pylori* exploits the versatile *cag* T4SS to tip the delicate STING signaling balance towards inflammatory responses that may stimulate carcinogenesis and enable chronic colonization of the gastric niche.

## RESULTS

### *H. pylori* provokes multiple DNA surveillance systems in a *cag* T4SS-dependent manner

Previous studies demonstrate the capacity of *H. pylori* to stimulate DNA-sensing pattern recognition receptors, including TLR9 (Rad et al., 2009; Varga et al., 2016b), raising the hypothesis that microbial nucleic acids are actively translocated into host cells. In agreement with previous reports (Rad et al., 2009; Varga et al., 2016b), *H. pylori* challenge of HEK293 reporter cell lines stably transfected with TLR9 demonstrated that *cag* T4SS activity is required for TLR9 stimulation (**Fig. 1A,B**). Consistent with the observation that CagA is not translocated into HEK293 cells (Kumar Pachathundikandi et al., 2011; Varga et al., 2016b), disruption of *cagA* did not diminish levels of *H. pylori*-induced TLR9 activation (**Fig. 1A and Fig. S1A**). To confirm that a functional *cag* T4SS is required to activate TLR9, we co-cultured TLR9 reporter cells with a *cagL* isogenic mutant or the corresponding genetically complemented strain. Whereas inactivation of *cagL* abrogated TLR9 activation, complementation in a heterologous chromosomal locus rescued TLR9 stimulation to levels indistinguishable from the parental WT strain (**Fig. 1B**). In concert with previous investigations, these data demonstrate that *cag* T4SS activity, but not CagA delivery, is required for robust TLR9 signaling.

**Figure 1.**
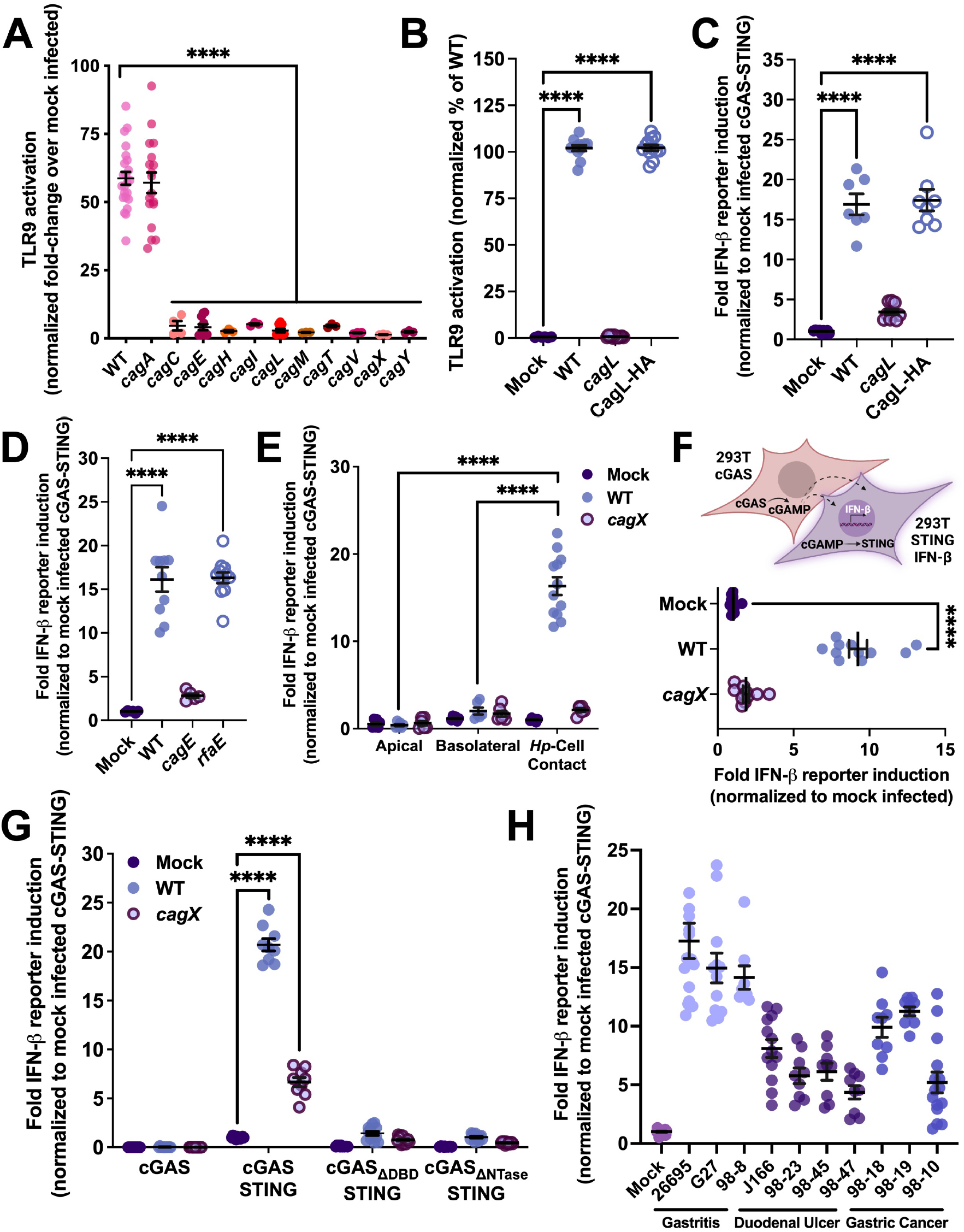
*H. pylori cag* T4SS activity stimulates multiple DNA surveillance systems. **A**. TLR9 activation induced by the indicated *H. pylori* 26695 isogenic mutant strain. Data are expressed as the normalized fold change over mock infected cells. **B**. TLR9 activation requires a functional *cag* T4SS. **C**. cGAS-STING signaling stimulated by the indicated strain. **D**. Induction of double-stranded DNA breaks in the host genome does not significantly contribute to *H. pylori*-induced cGAS-STING signaling. Graph depicts IFN-β reporter activity induced in cGAS-STING reporter cells by the indicated strain. **E**. Transwell cGAS-STING activation assays demonstrating the requirement for direct bacteria-host cell contact. **F**. STING transactivation assays providing evidence of intercellular cGAMP transfer. Schematic depicts the reporter cell line experimental strategy. **G**. IFN-β transcriptional reporter assays demonstrating the requirement of the cGAS DNA-binding domain (cGAS_ΔDBD_) and cGAS catalytic activity (cGAS_ΔNTase_) for *H. pylori*-induced cGAS-STING signaling. **H**. cGAS-STING signaling induced by the indicated *H. pylori* strain stratified by disease state (gastritis, duodenal ulcer, and gastric adenocarcinoma). In **A**-**G**, significance was determined by one-way ANOVA with Dunnett’s post-hoc correction for multiple comparisons to experimental controls. In all panels, ****, *p*<0.0001.

In addition to endosomal DNA pattern recognition receptors such as TLR9, gastric epithelial cells harbor cytosolic DNA surveillance proteins including the nucleotidyltransferase cyclic GMP-AMP (cGAMP) synthase (Cai et al., 2014; Diner et al., 2013; Sun et al., 2013; Wu et al., 2013). To test whether *H. pylori* DNA is trafficked into the host cell cytoplasm to activate cytosolic DNA surveillance sensors, we challenged 293T cells transfected with constructs to enable cGAS and STING expression and quantified levels of IFN-β promoter-driven luciferase produced in response to *H. pylori*. Compared to mock infected cells, WT *H. pylori* stimulated high levels of cGAS-STING signaling at 18 h post-infection. Similar to TLR9 activation assays, the *cagL* mutant was unable to stimulate robust cGAS-STING signaling, a phenotype that was restored by genetic complementation in a secondary chromosomal locus (**Fig. 1C**). In contrast to HEK293-hTLR9 cells, moderate levels of translocated CagA were detected in 293T cells co-cultured with WT *H. pylori* but not in corresponding *cagE*-challenged monolayers (**Fig. S1A**). Although *H. pylori* has the capacity to deliver CagA into 293T cells, disruption of *cagA* did not significantly alter levels of *cag* T4SS-dependent IFN-β promoter activity in cGAS-STING reporter cells (**Fig. S1B**). To exclude the possibility that cGAS-STING activation results from bacterial endocytosis or increased bacterial interaction with host cell surfaces, we performed gentamicin protection assays. In contrast to marked differences in cGAS-STING activation elicited by WT and *cagX*, equivalent levels of adherent and intracellular bacteria were recovered from 293T co-cultures, confirming that cGAS-STING signaling is not an artifact of non-specific bacterial internalization or spontaneous bacterial lysis (**Fig. S1C**).

Previous studies demonstrate that in addition to detecting invading microbial threats, cGAS senses and responds to DNA damage and genomic instability (Cai et al., 2014; Ke et al., 2022). *H. pylori cag* T4SS activity is directly linked to nuclear double-stranded DNA (dsDNA) breaks introduced in response to ALPK1/TIFA signaling stimulated by D-glycero-beta-D-manno-heptose 1,7-bisphosphate (HBP) or ADP-beta-D-manno-heptose (β-ADP-heptose) translocation (Bauer et al., 2020; Gall et al., 2017; Zimmermann et al., 2017). We therefore addressed the possibility that *cag* T4SS-dependent nuclear DNA damage stimulates cGAS-STING signaling elicited by WT *H. pylori*. Disruption of *rfaE*, the enzyme responsible for β-ADP-heptose production (Bauer et al., 2020; Gall et al., 2017; Stein et al., 2017; Zimmermann et al., 2017), did not impact the level of cGAS-STING signaling achieved by *H. pylori* (**Fig. 1D**), indicating that ALPK1/TIFA signaling-induced DNA damage does not significantly contribute to *H. pylori*-driven cGAS activation *in vitro*. In addition to ALPK1/TIFA-mediated DNA modifications (Bauer et al., 2020; Gall et al., 2017; Stein et al., 2017; Zimmermann et al., 2017), *H. pylori* has the capacity to induce production of DNA-damaging reactive oxygen species (ROS) by triggering NF-κB activation and additional mechanisms involving inducible nitric oxide synthase (iNOS) and associated inflammatory enzymes (Bauer et al., 2020; Kidane, 2018). In support of the observation that *cag* T4SS-induced dsDNA breaks are not a significant cGAS activating factor in the context of *H. pylori* infection, cGAS-STING activation was achieved by *H. pylori* co-cultured in the presence of the antioxidant N-acetyl-cysteine at concentrations that abrogate ROS production (Bauer et al., 2020) (**Fig. S1D**).

Following infection, damaged mitochondria release DNA (mtDNA) and other constituents into the cytosol to act as potent danger-associated molecular patterns (DAMPs) that engage TLR9 (Garcia-Martinez et al., 2016; Oka et al., 2012; Zhang et al., 2010) and cGAS-STING signaling axes to initiate type I IFN production (Rongvaux et al., 2014; West et al., 2015; White et al., 2014). To test the hypothesis that *H. pylori cag* T4SS activity modulates mitochondrial integrity resulting in the release of mtDNA and activation of cytosolic DNA-sensing PRRs, we assayed cGAS-STING activation in the presence of BAX/BAK macropore inhibitory peptides that prevent permeabilization of the mitochondrial outer membrane and herniation of mtDNA into the cytosol (McArthur et al., 2018; White et al., 2014). *H. pylori* induced similar levels of cGAS-STING activation in the presence of BAX inhibitory peptide or non-inhibitory peptide control co-cultures (**Fig. S1E**) suggesting that ruptured mitochondria are not the primary source of cGAS-activating DNA. Likewise, treatment of co-cultures with a mitochondria-targeted antioxidant did not significantly alter levels of cGAS-STING signaling (**Fig. S1D**), demonstrating that *H. pylori*-induced mtDNA damage is not a predominant DAMP within the context of *cag* T4SS-dependent cGAS activation. To exclude the possibility that cGAS-STING activation is dependent upon the import of released *H. pylori* DNA via host cell mechanisms, we quantified IFN-β promoter activity produced by cGAS-STING reporter cells that were physically separated from *H. pylori*. In support of the hypothesis that cGAS-STING activation requires direct *H. pylori*-host cell interaction, cGAS stimulation was achieved by *H. pylori* that were in direct contact with reporter cells, but not in samples in which *H. pylori* and reporter cells were physically separated by a 0.4 µM-pore polycarbonate insert (**Fig. 1E**). Collectively, these studies demonstrate that cGAS stimulation is a consequence of *cag* T4SS-dependent DNA delivery into the host cell cytosol.

To define the role of cGAS in *H. pylori*-driven IFN-β signaling, we next determined whether *cag* T4SS activity could transactivate STING through direct cGAMP transfer via gap junction-mediated diffusion. To assess cGAS signaling *in trans*, 293T cells transfected with cGAS constructs were co-cultured with 293T cells harboring STING and IFN-β reporter constructs, and co-cultures were challenged by *H. pylori* for 18 h. High levels of STING signaling were observed in cells challenged by WT *H. pylori*, but not a *cagX* mutant, suggesting that *cag* T4SS-dependent cGAS stimulation generates sufficient cGAMP for migration into bystander cells (**Fig. 1F**). To examine the requirement of cGAS protein domains in *H. pylori-*induced STING signaling, we next monitored IFN-β activation in 293T cells transfected with constructs to express either cGAS alone or cGAS variants in combination with STING. In comparison to cells transfected with only cGAS_WT_, WT *H. pylori* stimulated high levels of IFN-β transcription when cGAS_WT_ and STING were co-expressed (**Fig. 1G**). In contrast, *cag* T4SS-dependent IFN-β transcription was markedly reduced in 293T cells transfected with STING and cGAS variants lacking the DNA-binding domain (cGAS_ΔDBD_) or harboring point mutations within nucleotidyltransferase catalytic residues that abolish cGAMP production (cGAS_ΔNTase_) (**Fig. 1G**). Collectively, these results demonstrate that cGAS senses and responds to *cag* T4SS activity.

A previous report suggests that decreased STING signaling is associated with adverse outcomes in gastric cancer patients (Song et al., 2017). We therefore monitored cGAS-STING signaling induced by several *H. pylori* clinical isolates obtained from patients with gastric diseases of varying severity. In contrast to strains isolated from patients exhibiting gastritis, *H. pylori* isolated from individuals with duodenal ulcers or gastric cancer elicited lower levels of cGAS-STING signaling (**Fig. 1H**), suggesting that *H. pylori* may modulate the capacity to induce *cag* T4SS-dependent STING activation during chronic stomach colonization.

### Microbial DNA is delivered to the gastric epithelial cell cytoplasm via *cag* T4SS activity

We next assessed the consequence of *H. pylori* trans-kingdom DNA conjugation within the context of biologically-relevant interactions with gastric epithelial cells. To monitor DNA injection into gastric epithelial cells, AGS cells were challenged by either WT or the *cagX* isogenic mutant at a MOI of 50. After 6 h, gastric epithelial cell co-cultures were treated with DNaseI to remove extracellular DNA and eukaryotic cells were fractionated using digitonin to selectively permeabilize the plasma membrane, leaving the nuclear envelope and bacterial cells intact. PCR analysis of fractionated infected AGS cells revealed the presence of *H. pylori* DNA in cytoplasmic fractions (**Fig. 2A**) with significantly more bacterial DNA present in cytosolic fractions obtained from WT-challenged cells. When comparing the ratio of cytoplasmic bacterial DNA to cytoplasmic-localized mitochondrial DNA by qPCR, significantly more bacterial DNA was present in the cytosol of gastric epithelial cells infected by the WT strain compared to corresponding cells infected by the *cagX* mutant (**Fig. 2B**). Levels of mitochondrial DNA did not significantly differ in cytosolic extracts obtained from infected and uninfected cells (**Fig. 2A**). To exclude the possibility that cytosolic localization of *H. pylori* DNA resulted from non-specific bacterial lysis or endocytosis, we performed studies to analyze the level of adherent and internalized *H. pylori* in infected gastric epithelial cells. Using gentamicin-protection assays, we determined that similar levels of adherent and intracellular WT and *cagX* bacteria were recovered from AGS co-cultures (data not shown). Multiple cancer cell lines, including cell lines derived from gastric adenocarcinoma, continuously export low levels of extracellular cGAMP that serves as a potent immunotransmitter (Carozza et al., 2020). We thus measured levels of extracellular cGAMP secreted by AGS cells in response to *H. pylori* challenge. Consistent with the hypothesis that *cag* T4SS-mediated DNA delivery stimulates cytosolic cGAS and the subsequent production of cGAMP, the level of extracellular cGAMP was significantly higher in supernatants obtained from WT-challenged co-cultures compared to mock infected or corresponding *cagX*-challenged monolayers (**Fig. 2C**). Collectively, these results suggest that the presence of bacterial DNA in the host cell cytoplasm is a consequence of *cag* T4SS activity and leads to the production of cGAS-generated cGAMP.

**Figure 2.**
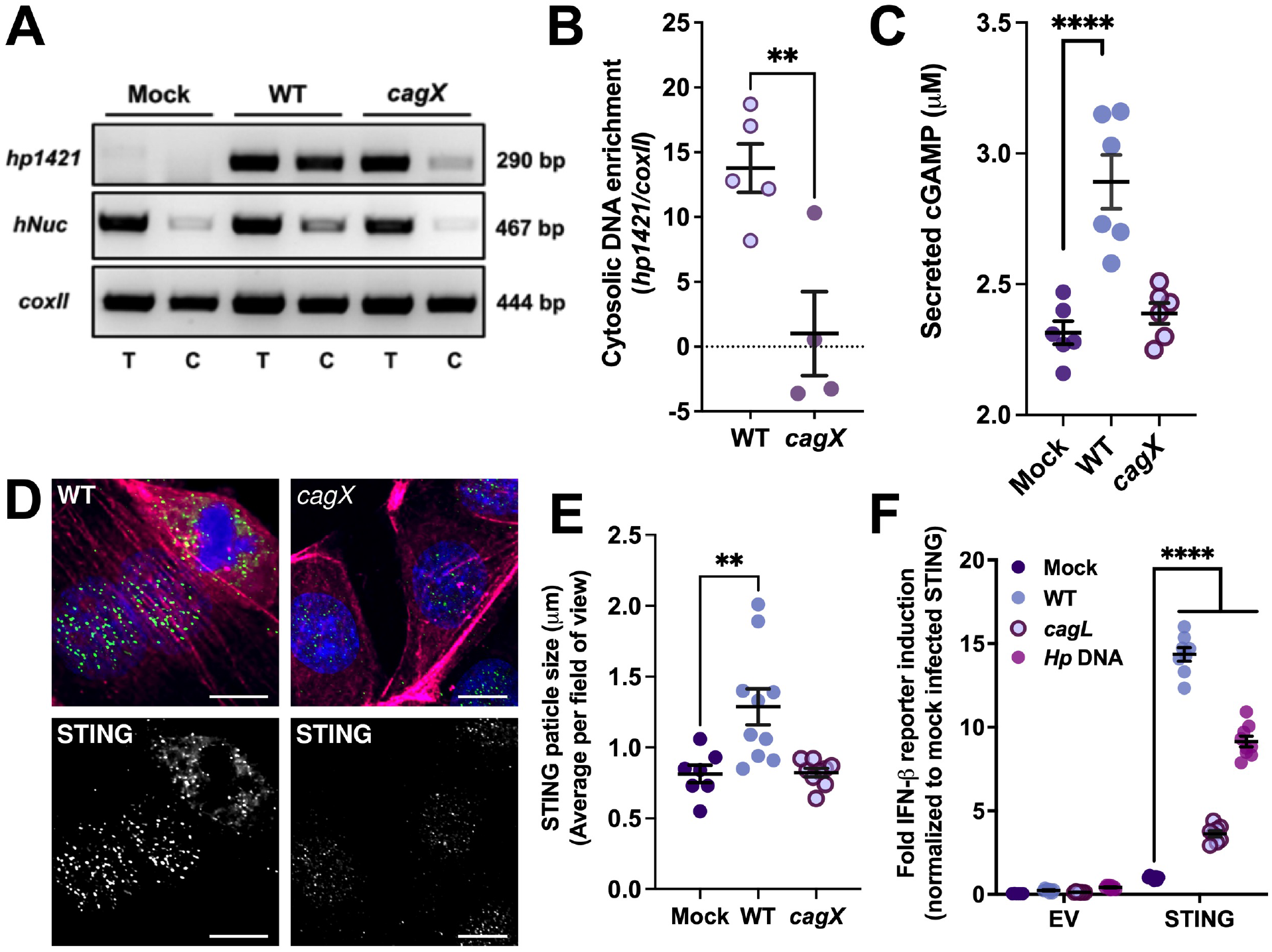
*H. pylori* DNA is delivered to the gastric epithelial cell cytoplasm in a *cag* T4SS-dependent manner. **A**. Representative PCR amplifications demonstrating the presence of chromosomal *H. pylori* (*hp1421*), nuclear genomic (*hNuc* (Fernandez-Moreno et al., 2016)), and mitochondrial (*coxII* (Fernandez-Moreno et al., 2016)) DNA fragments in fractionated cytoplasmic (C) and total (T) co-culture AGS cell extracts. **B**. qPCR analysis of *H. pylori* DNA enrichment in cytosolic fractions normalized to levels of cytosolic mitochondrial DNA. Results are representative of at least 4 biological replicate experiments. Significance was determined by unpaired, two-tailed t-test; **, *p*<0.01. **C**. Levels of extracellular cGAMP produced by AGS cells in response to *H. pylori*. **D**. Confocal microscopy analysis of perinuclear STING localization in *H. pylori*-challenged primary gastric epithelial cells at 6 h post-infection. Representative image of *n*=2 biological replicate experiments depicting STING (green), nuclei (blue), and actin (magenta) staining. Scale bar represents 20 µm. **E**. Quantitation of STING particle size in primary gastric epithelial cells challenged by the indicated *H. pylori* strain. Data represents the average STING particle size per field of view for mock infected (7 fields of view, *n*=90 cells); WT infected (11 fields of view, *n*=82 cells); and *cagX* infected (9 fields of view, *n*=59 cells) gastric epithelial cells. **F**. STING signaling induced by the indicated *H. pylori* strain or purified *H. pylori* chromosomal DNA in 293T reporter cells. Data is representative of a minimum of 3 biological replicate experiments. In **E*-*F**, significance was determined by one-way ANOVA with Dunnett’s post-hoc correction for multiple comparisons to experimental controls; ****, *p*<0.0001.

Although gastric adenocarcinoma cell lines (including AGS and MKN45) produce detectable levels of cGAS and other nucleic acid-sensing PRRs, STING expression is absent (Qiao et al., 2020). To determine whether translocated *H. pylori* DNA elicits cGAS-STING responses in normal gastric epithelia, we challenged primary adult gastric epithelial cells with *H. pylori* and monitored the formation of peri-nuclear STING polymers that aggregate in response to cGAMP binding. Compared to mock infected or *cagX*-challenged cells, WT *H. pylori* induced the formation of large STING aggregates that could be visualized by confocal microscopy (**Fig. 2D**). Consistent with activation of cGAS-STING signaling, quantification of the average STING polymer size in *H. pylori*-gastric cell co-cultures revealed significantly larger STING aggregates in WT-challenged cells (**Fig. 2E**), suggesting *cag* T4SS-dependent stimulation of cGAS surveillance. In addition to cGAS, epithelial cells harbor other nucleic acid reconnaissance systems that signal through STING. To determine whether *H. pylori* stimulates additional STING-dependent signaling pathways, we analyzed IFN-β activity in 293T-STING cells. Compared to control 293T cells harboring empty vector, *H. pylori* stimulated STING signaling in a *cag* T4SS-dependent manner (**Fig. 2F**). STING signaling was also elicited by transfection of purified, fragmented *H. pylori* chromosomal DNA into IFN-β reporter cells (**Fig. 2F**), suggesting that endogenous 293T cytosolic DNA-sensing pathways respond to *H. pylori* DNA. Together, these studies demonstrate that *H. pylori* delivers chromosomal DNA fragments to gastric epithelial cells via *cag* T4SS mechanisms and reveal that endogenous STING-dependent nucleic acid surveillance systems are activated by translocated *H. pylori* DNA.

### DNA is a specific *cag* T4SS nucleic acid substrate

We reasoned that in addition to DNA substrates, the *cag* T4SS may translocate RNA or DNA:RNA hybrids into gastric cells to stimulate STING-dependent signaling. Thus, we sought to characterize the innate inflammatory signature elicited by *H. pylori*. Primary adult gastric epithelial cells were mock infected or challenged by either WT or the corresponding *cagX* isogenic mutant prior to isolation of total RNA. To characterize epithelial innate inflammatory responses, gene expression patterns were analyzed at 6 h post-infection using the NanoString Host Response Panel. Compared to mock infected and *cagX*-challenged primary gastric epithelial cell co-cultures, *cag* T4SS activity induced a significant increase in transcripts associated with anti-microbial defense pathways and interferon signaling (**Fig. 3A and Fig. S2**). Hierarchical clustering performed on genes that were differentially expressed in response to *H. pylori* challenge revealed distinct clustering of mock infected and *cagX*-infected co-cultures in comparison to WT challenge (**Fig. S2C**). Comparison of transcripts differentially produced in response to WT versus *cagX H. pylori* identified 134 genes that were upregulated via *cag* T4SS activity, including genes encoding pro-inflammatory chemokines and cytokines (*e*.*g*., *CXCL10, CXCL8/IL-8, TNF, IL1B, CCL5*), IFN-*α*/*β* signaling (*e*.*g*., *ISG15, IRF1, IFI35, STAT1, IFNAR2, SAMHD1*), and IFN-γ immunoregulatory programs (*e*.*g*., *IFNGR2, TRIM5, GBP4, VCAM1*) (**Fig. 3A,B**). Consistent with the observed gene expression patterns, pathway analysis of genes induced by *cag* T4SS activity demonstrated a significant enrichment in transcripts associated with interferon-regulated nucleic acid reconnaissance programs (**Fig. 3C**). In addition to *cag* T4SS-dependent regulation of the DNA sensor ZBP1, we observed significantly increased transcript levels of several cytosolic RNA-sensing surveillance systems including *DDX58/RIG-I*, RIG-I-like Receptor *IFIH1/MDA5* (Chow et al., 2015; Dixit and Kagan, 2013), and 2′-5′-oligoadenylate synthetases *OAS1* and *OAS2* (Schwartz et al., 2020) in WT-infected primary gastric epithelial cells (**Fig. 3A,B**).

**Figure 3.**
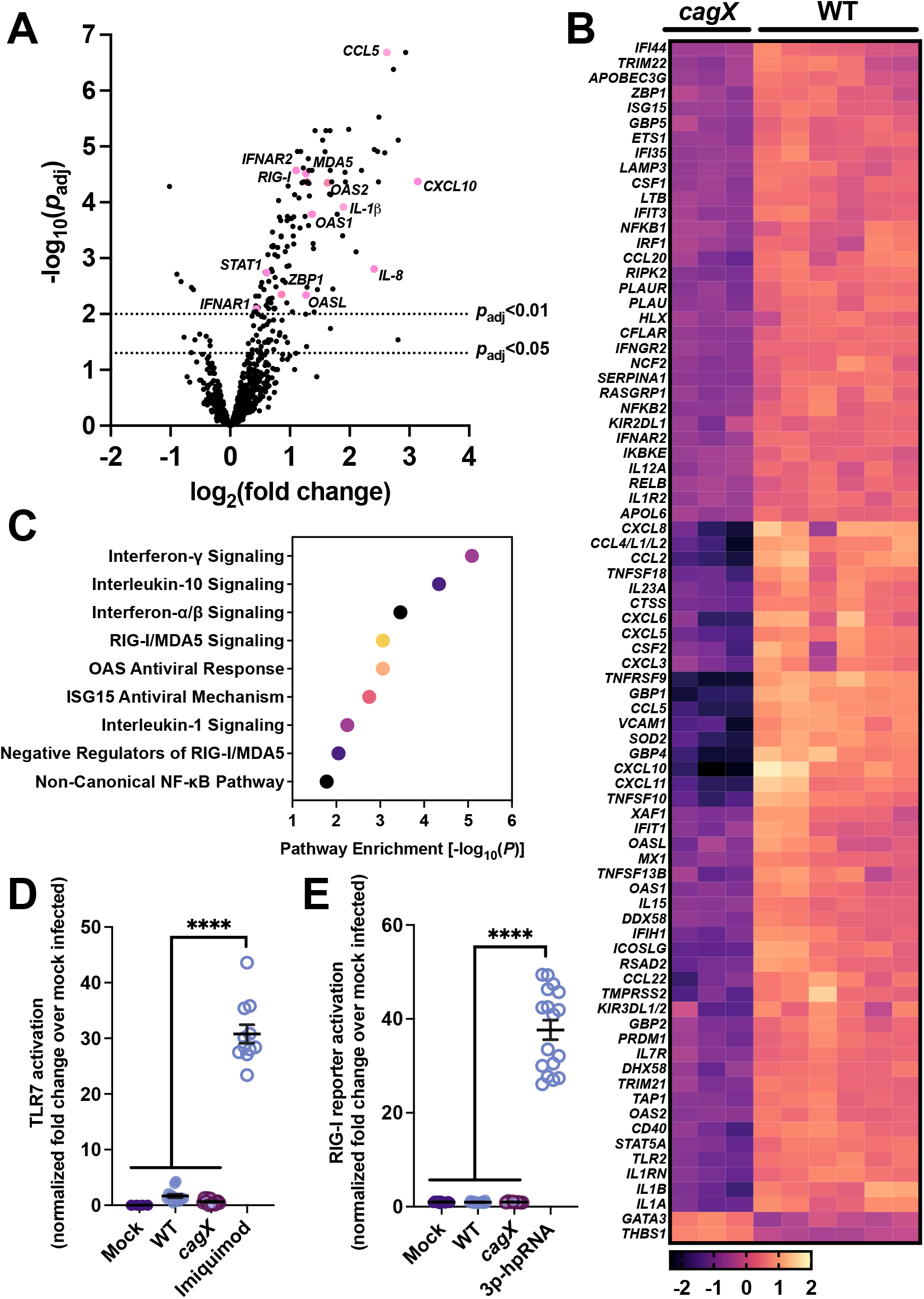
*H. pylori* regulates nucleic acid reconnaissance pathways via *cag* T4SS activity. **A**. Volcano plot depicting expression of immune-related genes in adult human primary epithelial cells challenged by *H. pylori* detected with the NanoString human host response panel. Graph represents the fold change and associated *p*-value of all differentially expressed genes in the panel for WT vs. *cagX* challenged cells. Dashed lines dashed lines demarcate genes meeting the threshold for significance (*p*_adj_<0.01 and *p*_adj_<0.05) after correction with the Benjamini–Hochberg procedure for controlling FDR. Selected genes encoding nucleic acid sensing pathways, interferon-responsive elements, and inflammatory cytokines/chemokines are labeled and indicated in pink. **B**. Heat map of differentially expressed genes depicted in a. Map depicts genes that were increased or decreased by 1.8-fold and an adjusted *p*-value <0.01. **C**. Pathway analysis of differentially expressed immune genes. Graph depicts the -log_10_ *p*-value for the indicated pathway. **D**. TLR7 activation levels induced by the indicated strain or pharmacological stimulus. **E**. Levels of RIG-I signaling stimulated by *H. pylori* or transfected RNA agonist. In **D** and **E**, significance was determined by one-way ANOVA with Dunnett’s post-hoc correction for multiple comparisons to experimental controls; ****, *p*<0.0001. Data is derived from *n*=1 NanoString analysis with gastric epithelial cell samples derived from *n*=2 biological replicate experiments (mock infected, *n*=3, WT infected *n*=6, and *cagX* infected *n*=3 samples analyzed).

A previous study reported the capacity of gastric mucosa-associated dendritic cells to sense and respond to purified *H. pylori* RNA through RIG-I and TLR7/8, leading to the production of type I IFN (Rad et al., 2009; Salama et al., 2013). However, whether the *cag* T4SS actively translocates RNA to stimulate TLR7/8 or RIG-I signaling axes has not been elucidated. We thus sought to determine whether DNA is a specific *cag* T4SS nucleic acid effector. Challenge of HEK293 reporter cell lines expressing either TLR7 (**Fig. 3D**) or RIG-1 (**Fig. 3E**) revealed that in contrast to robust activation of DNA-sensing PRRs, *H. pylori* was unable to activate either endosomal ssRNA (TLR7) or cytoplasmic ssRNA/dsRNA (RIG-I) sensors. Collectively, these data suggest that chromosomally-derived DNA is a specific *cag* T4SS substrate, and provide further evidence that *cag* T4SS-dependent DNA translocation stimulates STING-dependent signaling.

### *H. pylori cag* T4SS activity stimulates DNA immunosurveillance in bystander cells

Prior work demonstrates that foreign intracellular bacterial DNA can be delivered to adjacent cells via extracellular vesicles as a mechanism to amplify IFN signaling (Nandakumar et al., 2019). We therefore investigated whether translocated *H. pylori* DNA could be sorted into extracellular vesicles to enable paracrine-like DNA signaling by primary gastric epithelial cells. To test the hypothesis that infected gastric epithelial cells release extracellular vesicles containing *H. pylori* DNA, cell-free supernatants collected from infected primary gastric epithelial donor cells were used to challenge recipient TLR9 reporter cells. Compared to supernatants obtained from either mock infected or *cagX*-challenged co-cultures, supernatants harvested from WT-challenged gastric cells robustly activated TLR9 signaling (**Fig. 4A**). The capacity of infected cell supernatants to activate TLR9 was time dependent, as levels of TLR9 activation increased when reporter cells were treated with supernatants obtained from gastric epithelial cells co-cultured with *H. pylori* for 12 h compared to 6 h (**Fig. 4A**). To exclude the possibility that TLR9 activation resulted from contaminate DNA arising from cell lysis, we quantified the level of cell-free DNA in donor supernatants. TLR9 activation elicited by gastric cell supernatants was not correlated to the level of donor supernatant cell-free DNA (**Fig. 4B**), suggesting that the TLR9-stimulating agonist was enclosed within a host cell-derived delivery mechanism.

**Figure 4.**
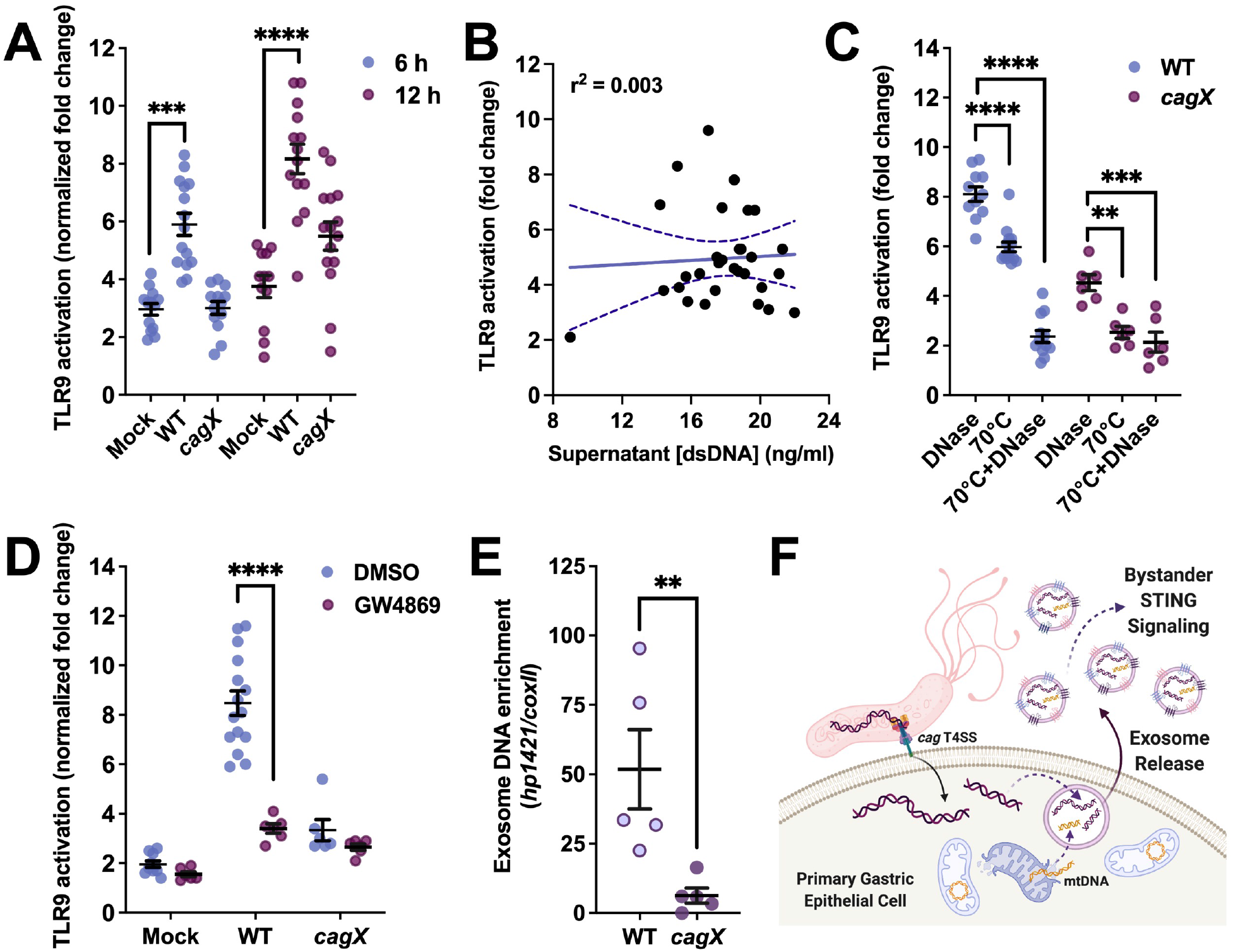
*H. pylori* effector DNA is packaged into exosomes to enable DNA pattern recognition receptor signaling in bystander cells. **A**. TLR9 stimulation induced by supernatants obtained from primary gastric epithelial cells challenged by *H. pylori* at the indicated time point post-infection. Graph depicts levels of TLR9 activation achieved by supernatants collected in a minimum of four biological replicate experiments. **B**. Linear regression analysis revealing no correlation between levels of TLR9 activation induced by gastric epithelial cell supernatants and the corresponding level of supernatant total cell-free DNA. **C**. Levels of TLR9 activation achieved by gastric cell supernatants obtained at 6 h post-infection and processed by the indicated conditions. **D**. Induction of TLR9 stimulation by gastric cell supernatant extracellular vesicles in gastric epithelial cell supernatants challenged by *H. pylori* in the presence or absence of GW4869 (10 μM). Significance was determined by unpaired, two-tailed t-test; ****, *p*<0.0001. **E**. qPCR analysis of *H. pylori* DNA enrichment in purified exosomes derived from primary gastric epithelial co-culture supernatants 6 h post-bacterial challenge by the indicated strain. Graph depicts the fold enrichment of *H. pylori* DNA (*hp1421*) over levels of mitochondrial DNA (*coxII*) in exosomes purified from supernatants collected from five biological replicate experiments. Significance was determined by unpaired, two-tailed t-test; **, *p*<0.01. **F**. Schematic representing a proposed model of translocated *H. pylori* DNA packaging and subsequent release of extracellular vesicles by primary gastric cells. In **A** and **C**, significance was determined by one-way ANOVA with Dunnett’s post-hoc correction for multiple comparisons to experimental controls; ****, *p*<0.0001, ***, *p*<0.001, and **, *p*<0.01.

To confirm the role of translocated DNA in transferrable nucleic acid immunosurveillance in bystander cells, donor gastric cell supernatants were treated with DNase alone or in combination with heat prior to co-culture with TLR9 reporter cells. Whereas DNase treatment had a negligible effect on TLR9 stimulating capacity, heat treatment modesty reduced TLR9 responses, which were further reduced when donor supernatants were treated with both heat and DNase prior to reporter cell challenge (**Fig. 4C**). These observations led to the hypothesis that translocated microbial DNA packaged within gastric cell-derived extracellular vesicles, such as exosomes, stimulates nucleic acid reconnaissance systems in bystander cells. To test whether exosome biogenesis is required for the delivery of translocated DNA to uninfected cells, we challenged recipient TLR9 reporter cells with donor supernatants obtained from infected primary gastric epithelial cells cultured in the presence or absence of a neutral sphingomyelinase (nSmase2) inhibitor that prevents exosome release. Compared to untreated and mock infected cells, nSmase2 inhibition led to a significant reduction in TLR9 activation levels elicited by WT-challenged donor cell supernatants (**Fig. 4D**), supporting a role of exosomes in stimulating DNA-sensing pathways in bystander cells. To further investigate whether translocated *H. pylori* DNA was packaged within exosomes released from infected gastric epithelial cells, we isolated CD9, CD63, and CD81-positive exosomes from cell culture supernatants via immunopurification. Analysis by qPCR revealed that exosomes purified from WT-challenged gastric cell supernatants were significantly enriched in *H. pylori* DNA compared to corresponding exosomes obtained from *cagX*-challenged co-cultures (**Fig. 4E**), suggesting that foreign bacterial DNA is sorted into exosomes for paracrine-like DAMP signal amplification (**Fig. 4F**).

### Random chromosomal fragments are delivered to target cells via *cag* T4SS-dependent mechanisms

To identify whether a specific DNA sequence is excised and transferred to host cells by *cag* T4SS activity, we purified cGAS-DNA complexes using modified ChIP-seq workflows to capture bacterial DNA that physically binds and activates cGAS. 293T-cGAS cells were challenged by *H. pylori* for 6 h, followed by chemical cross-linking and immunopurification of cGAS. Deep sequencing analysis confirmed that co-purifying *H. pylori* DNA isolated from WT and *cagX*-infected cells mapped across the entire *H. pylori* chromosome, with significantly more bacterial DNA reads associated with cGAS purifications obtained from WT-challenged monolayers (**Fig. 5A**). Normalized sequencing reads and differential peak calling approaches identified more than three hundred *H. pylori* chromosomal regions specifically associated with cGAS purified from WT-infected cells compared to eight bacterial DNA peaks associated with corresponding preparations obtained from *cagX* co-cultures (*P*=0.01). When enriched peak centers were mapped to the coordinate position across the *H. pylori* chromosome, differentially enriched peaks heavily clustered around the *oriC* region, with few peaks mapping to the chromosomal region diametrically opposed to *oriC* (**Fig. 5B**), suggesting that DNA translocation is linked to bi-directional DNA replication.

**Figure 5.**
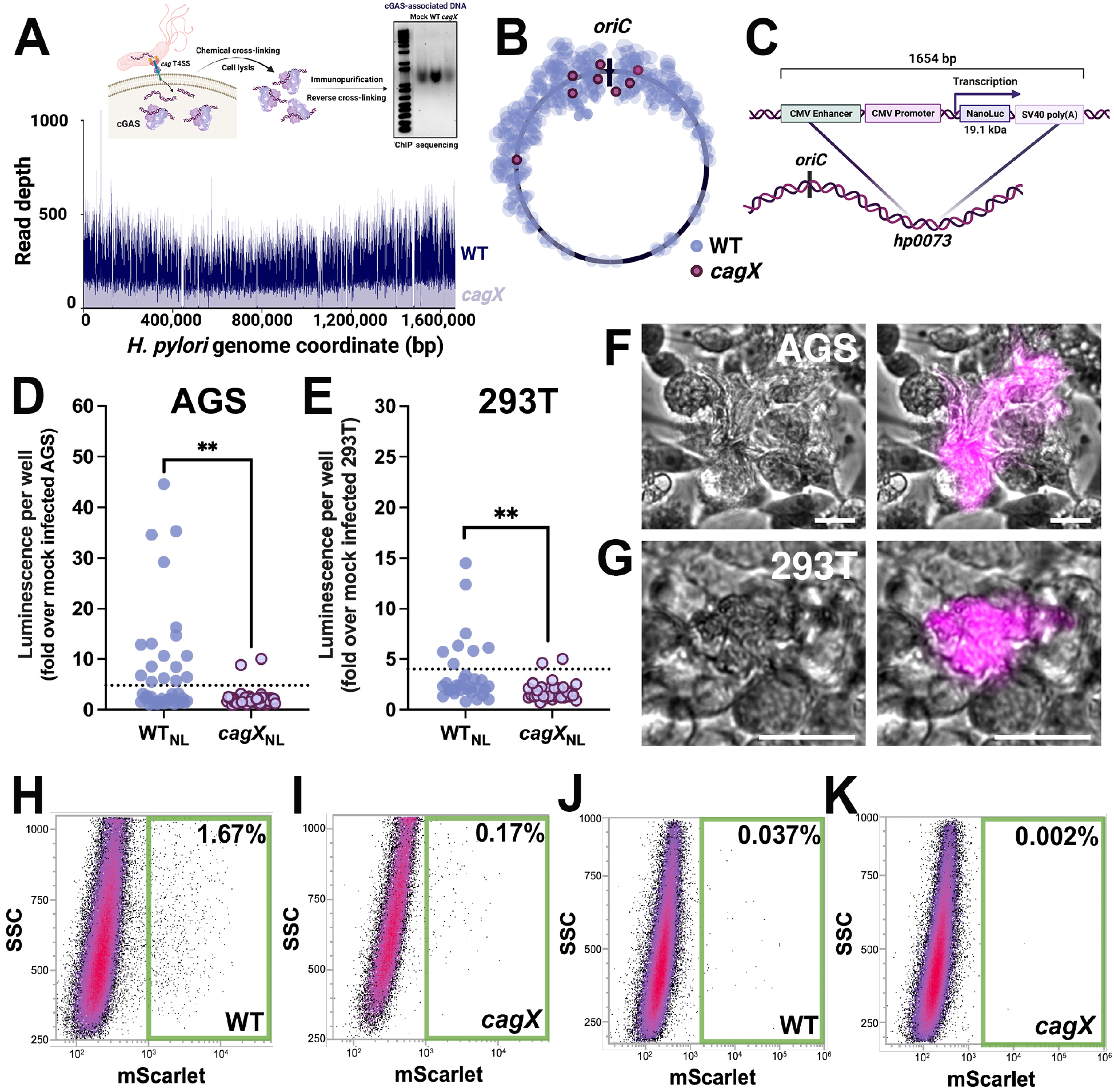
Chromosomally-derived DNA is translocated into target cells via *cag* T4SS mechanisms. **A**. Merged tracks of mapped *H. pylori* DNA reads obtained from infected 293T cGAS cells. Graph depicts sequencing read depth versus nucleotide position in the *H. pylori* 26695 genome. Schematic illustrates the experimental workflow for cGAS ‘ChIP-seq’ studies. **B**. DNA reads co-purified with cGAS were normalized to reads obtained from mock infected cells, and peak calling was used to identify regions of bacterial DNA that were enriched with cGAS immunopurification. Dots depict individual peaks and the corresponding peak center on the *H. pylori* 26695 chromosome obtained from WT (purple dots) and *cagX* (maroon dots) challenged co-cultures. **C**. Schematic of eukaryotic-optimized nanoluciferase expression constructs inserted into the *hp0073* locus. Nanoluciferase constructs were inserted frameshifted in the opposite orientation of the native operon transcription. **D**,**E**. Nanoluciferase bioluminescence produced by AGS (**D**) and 293T (**E**) cells challenged by the indicated strain at 24 h post-infection. Data represent a minimum of four biological replicate experiments. Significance was determined by unpaired, two-tailed t-test; **, *p*<0.01. **F**,**G**. Live cell phase contrast and fluorescence microscopy analysis of AGS (**F**) and 293T (**G**) cells challenged by WT *H. pylori* harboring LifeAct-mScarlet expression constructs at 24 h post-infection. Images are representative of *n*=2 biological replicate experiments. **H-K**. Flow cytometry analysis of AGS (**H** and **I**) or 293T cells (**J** and **K**) challenged by WT[mScarlet] or *cagX*[mScarlet] at 18 h post-infection. Green boxes indicate gating of mScarlet positive cells. Data is representative of *n*=2 biological replicate experiments.

To test the hypothesis that chromosomal regions near *oriC* are translocated to gastric epithelial cells at a high frequency, we cloned a CMV promoter-driven monomeric nanoluciferase construct into the *ureA* locus (*hp0073*) adjacent to *oriC* (**Fig. 5C**). Gastric epithelial cells and 293T cells were challenged by WT or *cagX H. pylori* strains harboring the eukaryotic-optimized nanolucifersase expression construct (WT_NL_ and *cagX*_NL_, respectively), and luciferase activity in infected cell lysates was monitored by bioluminescence. Compared to mock infected and *cagX*_NL_-infected cells, AGS and 293T cells challenged by WT_NL_ produced high levels of bioluminescence (**Fig. 5D** and **5E**), indicating that nanoluciferase production by epithelial cells is linked to *cag* T4SS-dependent DNA translocation.

To confirm that targeted DNA fragments proximal to *oriC* can be transferred to host cells via cag T4SS mechanisms, we replaced the nanoluciferase gene with a eukaryotic-optimized construct designed to express monomeric mScarlet tethered to the LifeAct N-terminal peptide to enable live cell visualization of F-actin. Microscopy analysis of AGS (**Fig. 5F**) and 293T (**Fig. 5G**) monolayers challenged by WT_mScarlet_ revealed numerous mScarlet-positive cells compared to corresponding *cagX*_mScarlet_-infected co-cultures. Flow cytometry analysis confirmed our observation that significantly more mScarlet-positive epithelial cells were generated when monolayers were challenged by WT_mScarlet_ compared to T4SS-deficient controls (**Fig. 5H-K**). Together, these data provide direct evidence that targeted chromosomal fragments greater than 1.5 kb in length are excised and delivered to host cells via *cag* T4SS-dependent mechanisms.

### *H. pylori* trans-kingdom conjugation is mechanistically coupled to chromosomal decatenation

We next sought to understand the mechanism by which effector DNA is coupled to the *cag* T4SS for transfer to target host cells. The observation that fragments of translocated bacterial DNA map predominantly to the *oriC* region led to the hypothesis that *H. pylori* trans-kingdom conjugation is linked to chromosomal replication. In support of this hypothesis, *cag* T4SS-dependent TLR9 activation was significantly impaired in the presence of the DNA gyrase inhibitor ciprofloxacin in a dose-dependent manner (**Fig. 6A**). To analyze the contribution of DNA segregation proteins to coupling transfer DNA to the *cag* T4SS apparatus, we employed targeted mutagenesis to delete individual genes known to be involved in DNA partitioning. Whereas isogeneic mutants deficient in genes encoding the DNA partitioning proteins ParA or ParB induced WT levels of *cag* T4SS-depenedent TLR9 activation, isogenic mutants lacking genes encoding the DNA translocase *ftsK* (*hp1090*) or the recombinase *xerH* (*hp0675*) exhibited marked defects in TLR9 stimulation (**Fig. 6B**), suggesting that chromosomal dimer resolution is required for transfer DNA coupling to the *cag* T4SS apparatus. TLR9 stimulation defects exhibited by the *ftsK* mutant could be rescued by genetic complementation with full length FtsK, but not a truncated FtsK harboring only the translocase DNA binding domain (FtsK-γ), which is required for interaction with XerH and XerH-mediated DNA recombination (Debowski et al., 2012) (**Fig. 6B**). Loss of *xerH* or *ftsK* was not associated with defects in *cag* T4SS-dependent induction of IL-8 synthesis by gastric epithelial cells (**Fig. 6C**), indicating that chromosomal segregation defects specifically impair *cag* T4SS phenotypes associated with DNA translocation. Consistent with the observed defects in TLR9 activation, *ftsK* mutants stimulated significantly reduced levels of cGAS-STING signaling compared to WT and the corresponding FtsK complemented strain (**Fig. 6D**).

**Figure 6.**
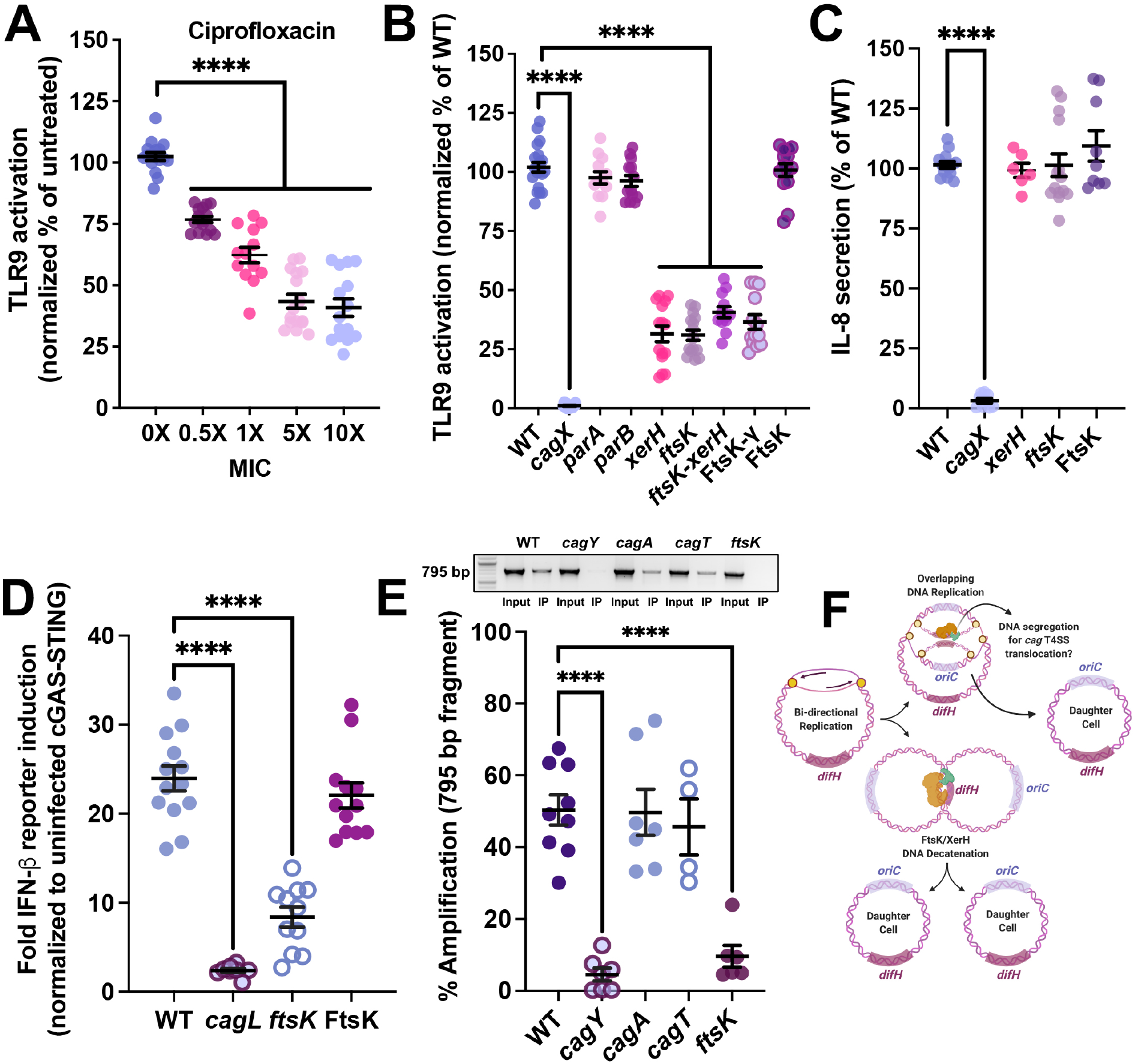
DNA translocation is mechanistically coupled to chromosomal replication and replichore decatenation. **A**. TLR9 activation induced by WT *H. pylori* in the presence of the indicated ciprofloxacin minimum inhibitory concentration (1X MIC = 0.125 µg/ml). Data are expressed as a percent of TLR9 stimulation achieved by WT in mock treated wells. **B**. Levels of TLR9 activation induced by the indicated isogenic mutant. Data are expressed as a percent of TLR9 stimulation achieved by the parental WT strain. **C**. IL-8 secreted by AGS cells challenged by the indicated *H. pylori* strain at 4.5 h post-infection. Data are expressed as a percent of IL-8 levels stimulated by the corresponding WT strain. **D**. cGAS-STING signaling induced by the indicted strain. Graph depicts the results of *n*=3 biological replicate experiments. **E**. Transfer DNA immunopurification assays demonstrating the presence of *H. pylori* chromosomal DNA fragments within the *cag* T4SS apparatus. Graph depicts the amplification efficiency of a 795 bp fragment in transfer DNA assay preparations purified from the indicated strain. Amplification efficiency of the immunopurification (IP) samples are expressed as the percent of the input DNA from at least four biological replicate experiments. Inset depicts representative PCR amplifications obtained from input and IP samples prepared from the indicated strain. **F**. Proposed model of DNA cargo segregation for *cag* T4SS-dependent delivery to host cells. We hypothesize that FtsK-XerH complexes mediate the rare excision of DNA arising from overlapping rounds of chromosomal replication for subsequent coupling to the *cag* T4SS apparatus for delivery to host cells. In **A**-**E**, significance was determined by one-way ANOVA with Dunnett’s post-hoc correction for multiple comparisons to experimental controls; ****, *p*<0.0001 in all panels.

To investigate the role of chromosomal segregation in loading effector DNA into the *cag* T4SS apparatus, we developed a ‘transfer DNA’ immunopurification assay to capture chromosomal fragments contained within the *cag* T4SS translocation channel. We reasoned that effector DNA trafficked into the *cag* T4SS apparatus would physically interact with components of the outer membrane complex that comprise the secretion chamber (Chung et al., 2019; Frick-Cheng et al., 2016). To test this hypothesis, we treated *H. pylori* with formaldehyde to chemically cross-link DNA-protein complexes, lysed the cells by sonication, and purified the *cag* T4SS core complex via immunopurification targeting CagY, which forms the central cap-like structure within the inner ring of the outer membrane complex (Chung et al., 2019). PCR analysis targeting a 795 bp chromosomal fragment revealed the presence of DNA in reverse cross-linked CagY complexes purified from WT but not in mock preparations obtained from *cagY* isogenic mutants (**Fig. 6E**). DNA co-purification with CagY was not dependent on the presence of either the effector protein CagA or CagT, a component localized to the periphery of the outer membrane ring complex (Chung et al., 2019; Frick-Cheng et al., 2016); however, DNA loading into the *cag* T4SS apparatus was significantly impaired in the absence of FtsK (**Fig. 6E**). Together, these data demonstrate that effector DNA is loaded into the *cag* T4SS machinery prior to encountering host cells to establish a ‘ready-to-fire’ nanomachine and demonstrate that transfer DNA loading is mechanistically coupled to chromosomal replication and replichore decatenation. We thus propose a model in which FtsK-XerH complexes resolve imbalanced replichores arising from overlapping rounds of chromosomal replication, resulting in the rare excision of DNA that is subsequently shuttled through the *cag* T4SS apparatus via unknown mechanisms (**Fig. 6F**).

## DISCUSSION

STING-dependent immunosurveillance plays a critical role in maintaining gastric mucosal homeostasis and regulating inflammatory responses with carcinogenic potential (Ke et al., 2022). STING signaling has been implicated in the development and progression of stomach cancer, but whether STING activation promotes or restricts gastric carcinogenesis remains unresolved. For example, a previous study demonstrated that STING downregulation in primary gastric tumors was associated with increased malignancy and the progression of gastric cancer (Song et al., 2017), while a conflicting report determined that high STING expression in malignant tissues and tumor-associated macrophages was predictive of poor prognosis in stomach cancer patients (Miao et al., 2020). Thus, the molecular role of cGAS-STING signaling in chronic gastric inflammation and pre-malignant lesion development remains unresolved. Consistent with the observation that chronic *H. pylori* colonization elicits STING signaling in the murine gastric mucosa (Song et al., 2017), our study establishes a critical role of *cag* T4SS-mediated trans-kingdom DNA conjugation in stimulating STING-dependent outcomes that underscore infection-associated carcinogenesis.

In the context of bacterial infection, type I IFN responses have been associated with both protective and detrimental outcomes (Boxx and Cheng, 2016; Peignier and Parker, 2021). Previous work demonstrates that the *H. pylori cag* T4SS-dependent delivery of muropeptide fragments activates Nod1 sensing and IRF7-mediated type I IFN responses in epithelial cells that restrict bacterial proliferation via CXCL10-mediated immune cell recruitment (Viala et al., 2004; Watanabe et al., 2010). Our work demonstrates that during acute infection, CXCL10 is highly upregulated by *cag* T4SS activity in primary gastric epithelial cells (**Fig. 3**). In addition to increased transcript levels associated with interferon stimulated genes, we observed augmented production of transcripts associated with *SMHD1* [a suppressor of type I IFN and NF-*κ*B signaling (Chen et al., 2018)], *IFI35* [a negative regulator of RIG-I signaling (Das et al., 2014)], and *ISG15* [an IFN-α/β-inducible ubiquitin-like modifier that is a key negative regulator of IFN-α/β immunity (Zhang et al., 2015)] in WT-infected primary gastric cells, suggesting that *H. pylori* has evolved mechanisms to counteract or suppress nucleic acid signaling pathways induced by *cag* T4SS activity. We speculate that similar to evasion strategies employed by numerous viruses, *H. pylori* counteracts type I IFN responses through the *cag* T4SS-dependent translocation of as yet unidentified, evolutionarily-conserved protein effectors that target and neutralize components of DNA-sensing or STING signaling pathways. Alternatively, pro-inflammatory STING signaling may balance immunosuppressive responses stimulated by *cag* T4SS-dependent TLR9 activation (Varga et al., 2016a) as a mechanism to sustain gastric homeostasis during acute colonization.

Epithelial DNA damage, genomic alterations, and chromosomal instability are hallmarks of *H. pylori*-induced gastric cancer (Chaturvedi et al., 2014). While *H. pylori cag* T4SS activity has been shown to induce the formation of both double-stranded DNA breaks (DSBs) and micronuclei that potentially contribute to cGAS-STING signaling in gastric epithelial cells (Bauer et al., 2020; Hanada et al., 2014; Koeppel et al., 2015), our data suggest that translocated chromosomal DNA fragments serve as the predominant cytosolic cGAS trigger in the context of *H. pylori* infection (**Fig. 1** and **Fig. S1**). In support of our data indicating that translocated chromosomal fragments are the key agonist that alerts cytosolic nucleic acid surveillance, we found that *H. pylori* effector DNA is packaged into exosomes released by gastric epithelial cells to enable paracrine-like signal amplification (**Fig. 4**). Consistent with a previous study demonstrating that intracellular *Listeria monocytogenes* infection stimulates the production of extracellular vesicles with IFN-inducing potential (Nandakumar et al., 2019), the delivery of pathogen-derived DNA via exosome release may represent a conserved mechanism by which epithelial cells potentiate nucleic acid surveillance-dependent danger signaling to uninfected tissues. We propose that *H. pylori* has evolved to exploit this innate defense mechanism to shape a tolerogenic immune response that favors persistent gastric colonization (Varga et al., 2016a).

Mechanistically, our data demonstrate that *cag* T4SS-dependent DNA delivery is coupled to chromosomal replication and replichore decatenation (**Fig. 5** and **Fig. 6**). Similar to *Neisseria gonorrhoeae* ssDNA secretion mechanisms (Callaghan et al., 2017), chromosomal partitioning influences *cag* T4SS-mediated DNA translocation. Indeed, our studies reveal that effector DNA coupling to the *cag* T4SS apparatus requires both FtsK membrane tethering and DNA translocase activity (**Fig. 6**). In support of the critical role of replichore decatenation in *cag* T4SS-dependent DNA translocation, *H. pylori* lacking the DNA recombinase XerH, which requires direct interaction with FtsK to resolve chromosome dimers and catenated chromosomes (Bebel et al., 2016; Debowski et al., 2012), exhibited significantly reduced levels of TLR9 stimulation. Our studies suggest that while *xerH* mutants harbor increased DNA content per cell compared to the parental WT strain (Debowski et al., 2012), *cag* T4SS DNA translocation phenotypes are significantly impaired when chromosomal DNA topological isomers cannot be efficiently resolved (**Fig. 6**). Additionally, our findings establish that DNA effector molecules are pre-loaded into the *cag* T4SS apparatus to establish a ‘ready-to-fire’ nanomachine that can be rapidly deployed during acute infection, and demonstrate that FtsK is required for partitioning DNA substrates into the *cag* T4SS (**Fig. 6**). Although *H. pylori* harbors several orthologs to paradigmatic DNA conjugation systems, endonucleases, phage integrases, and canonical *A. tumefaciens* VirD2 relaxases, the mechanism by which chromosomally-derived effector DNA is precisely excised and shuttled through the *cag* T4SS while maintaining faithful genome inheritance remains unresolved. Thus, further studies are warranted to explore the intriguing possibility that the *cag* T4SS has co-opted orphaned components of remnant *tfs3* and *tfs4* conjugation systems to enable trans-kingdom DNA delivery (Fernandez-Gonzalez and Backert, 2014).

Our results provide the first direct evidence that *H. pylori* delivers immunostimulatory chromosomal DNA fragments into target epithelial cells and demonstrate that translocated effector DNA physically binds and activates cytosolic DNA-sensing reconnaissance systems. The observation that chromosomal fragments encompassing full-length eukaryotic genes can be excised and directed to the *cag* T4SS for delivery into target cells underscores the potential to exploit *H. pylori* trans-kingdom conjugation for targeted DNA cargo delivery. Notably, our work provides the essential experimental framework required to harness the *cag* T4SS as a mucosal-targeted DNA vaccine or therapeutic delivery device that can be pharmacologically controlled *in vivo*. Studies to determine the maximal DNA fragment length that can be efficiently transported to gastric cells via *cag* T4SS mechanisms are currently underway.

Our work highlights the importance of understanding host-pathogen conflicts that engage cellular PRR signaling axes to drive chronic inflammation and stimulate the development of infection-related malignancies. While we propose that translocated microbial DNA plays a critical and underappreciated role in the development of inflammation-associated carcinogenesis, further investigations are required to understand STING-dependent outcomes within the context of gastric cancer. Our findings identify a central role of *cag* T4SS activity in eliciting cGAS-STING immunosurveillance in the gastric epithelium; however, additional studies are necessary to understand how STING activation shapes the gastric immune landscape to enable persistent *H. pylori* colonization and to elucidate whether STING signaling influences the development of pre-malignant lesions. Finally, our work more broadly points towards therapeutic STING modulation as a promising intervention strategy to reduce the incidence and severity of infection-associated malignancies that arise as a consequence of chronic inflammation.

## ACKNOWLEDGEMENTS

This work was funded by the NIH (P20 GM130456 and P30 GM110787 to CLS) and academic developments funds provided by the University of Kentucky (to CLS).

## METHODS

### Bacterial strains and culture conditions

*Helicobacter pylori* strain 26695, strain G27, isogenic derivatives, and clinical isolates were propagated on trypticase soy agar plates supplemented with 5% sheep blood (BD) as previously described (Shaffer et al., 2011). Overnight cultures of *H. pylori* were grown in Brucella broth supplemented with 5% fetal bovine serum (FBS) at 37°C with 5% CO_2_. For cloning purposes, *E. coli* DH5α (New England Biolabs) were grown on lysogeny broth (LB) agar plates or in LB liquid medium with appropriate antibiotics required for plasmid maintenance.

### Human cell culture

The AGS human gastric adenocarcinoma epithelial cell line (ATCC CRL-1739) was cultured in RPMI 1640 medium supplemented with 10% FBS, 2 mM L-glutamine, and 10 mM HEPES in the presence of 5% CO_2_ at 37°C. Primary adult human gastric epithelial cells (Cell Biologics H-6039) were grown in human epithelial cell medium supplemented with ITS, EGF, hydrocortisone, L-glutamine, and 5% FBS (Cell Biologics H6621) in collagen-coated cell culture flasks at 5% CO_2_ and 37°C. Prior to assays, wells of tissue culture plates were coated in collagen following manufacturer’s protocol. HEK293-hTLR9 (Invivogen hkb-htlr9), the corresponding parental HEK293 null1 (InvivoGen hkb-null1), HEK293-hTLR7 (Invivogen hkb-htlr7), HEK-Lucia™ hRIG-I (Invivogen hkl-hrigi), and 293T (ATCC CRL-3216) were grown in DMEM supplemented with 10% heat-inactivated FBS and 1X GlutaMAX (Life Technologies) in the presence of 5% CO_2_ at 37°C. For cell treatments prior to bacterial challenge, the following compounds were added at the indicated final concentration one hour prior to bacterial challenge: MitoTempo (Sigma-Aldrich, 100 μM); *N-*acetylcysteine (Sigma-Aldrich, 1 mM); BAX-inhibiting peptide, negative control (Sigma-Aldrich, 200 μM); BAX-inhibiting peptide V5 (Sigma-Aldrich, 200 μM); or sphingomyelinase inhibitor GW4869 (Sigma-Aldrich, 10 μM).

### *H. pylori* mutagenesis

Isogenic mutant derivatives of *H. pylori* 26695 and G27 were generated essentially as previously described (Johnson et al., 2014; Shaffer et al., 2011; Varga et al., 2021). Briefly, *H. pylori* was transformed with a suicide plasmid in which the coding region of the target gene was replaced by either a kanamycin resistance cassette or a chloramphenicol resistance cassette and homologous flanking DNA sequences 500 base pairs (bp) up-and downstream of the target locus. Colonies resistant to kanamycin (12.5 μg/ml) or chloramphenicol (10 μg/ml) were PCR-verified to confirm insertion of the resistance cassette into the appropriate locus. To complement mutants in *cis* at the *ureA* locus, plasmids derived from pAD1 (Shaffer et al., 2011) were constructed to express either the native gene or a hemagglutinin (HA) epitope-tagged protein as previously described (Shaffer et al., 2011; Varga et al., 2021). Plasmid sequences were confirmed by sequencing, and constructs were used to transform isogenic mutant strains. Insertion of complementation constructs into the *ureA* locus was confirmed by PCR amplification and anti-HA immunoblotting, when appropriate.

To construct *H. pylori* strains harboring NanoLuc luciferase expression constructs, a 1654 bp fragment encompassing the CMV enhancer element, CMV promoter, NanoLuc luciferase gene, and SV40 late poly(A) signal was amplified from pNL1.1.CMV[Nluc/CMV] (19.1 kDa NanoLuc protein, Promega). The PCR product was blunt end ligated to pAD1 digested with EcoRV. Ligation insertion was confirmed by PCR and DNA sequencing. Clones in which the NanoLuc luciferase construct was inserted in the reverse orientation relative to *ureA* transcription were selected for transformation into WT and *cagX H. pylori* 26695 and WT G27 strains. Integrations into the *H. pylori* chromosome were confirmed by PCR. The G27 isogenic *cagE* mutant was generated by replacement of the *cagE* coding region with a kanamycin resistance cassette as previously described (Shaffer et al., 2011; Varga et al., 2021). *H. pylori* strains harboring monomeric LifeAct-mScarlet (Bindels et al., 2017) expression constructs were generated using the same mutagenesis strategy, with the exception that a 1872 bp region of pLifeAct_mScarlet-i_N1 (Bindels et al., 2017) (26.4 kDa LifeAct-mScarlet protein, Addgene) encompassing the CMV enhancer element, CMV promoter, LifeAct-mScarlet gene, and SV40 late poly(A) signal was amplified was cloned into pAD1 via blunt end ligation. Clones in which the LifeAct mScarlet construct was inserted in the reverse orientation relative to *ureA* transcription were selected for transformation into *H. pylori* 26695 and G27 strains. Integrations into the *ureA* locus were confirmed by PCR.

### cGAS-STING reporter assays

293T cells seeded into 24-well plates at a density of approximately 1.5 × 10^5^ cells per well were transfected using Lipofectamine 2000 (Life Technologies) complexed with a combination of plasmids pUNO1-hSTING-R232 (Invivogen puno1-hStingWT), pUNO1-hcGAS-HA3X (Invivogen pUNO1-HA-hcGAS), cGAS derivatives (pUNO1-hcGAS-AA, Invivogen pUNO1-hcGAS-AA; pcDNA3.1-cGAS_ΔDBD_, Addgene 102606), pRL-SV40P (Addgene 27163), IFN-Beta-pGL3 (Addgene 102597), or empty vector pcDNA3.1 (Life Technologies V79020) to ensure equivalent DNA concentrations according to the manufacturer’s protocol. At 16 h post-transfection, cells were challenged by mid-log phase WT *H. pylori*, corresponding *cag* isogenic mutants, or *cag* T4SS+ clinical isolates at a multiplicity of infection (MOI) of 20 bacterial cells per 293T cell. Alternatively, STING reporter cells were transfected with purified *H. pylori* genomic DNA (500 ng/well) at 16 h post-initial transfection of STING and IFN-β reporter plasmids. For cGAS-STING time course assays, *H. pylori* were added to reporter cell monolayers at a MOI of 20, and culture plates were centrifuged at 1,800 x g for 1 min to synchronize infections. At the indicated time point, gentamicin was added at a final concentration of 100 µg/ml.

At 24 h post-infection, cell supernatants were removed, and adherent cells were lysed in luciferase assay passive lysis buffer (Pierce). Luciferase luminescence generated by 20 µl cell lysate per well in a 96-well plate format was recorded with a microplate reader (BioTek Synergy H1) using the Dual-Luciferase Reporter Assay System (Pierce). Firefly luciferase luminescence (D-Luciferin, IFN-Beta-pGL3) was normalized to Renilla luciferase luminescence (coelenterzine, pRL-SV40P) for each well, and IFN-β transcriptional reporter values were expressed as the normalized fold change over mock-infected wells. A minimum of three biological replicate experiments were performed in quadruplicate for each assay.

To evaluate the requirement of direct bacteria-host cell contact for cGAS-STING signaling, 293T cells were seeded directly into the well of a 24-well plate or into a 0.4 µm pore size transwell insert at approximately 5 × 10^4^ cells per transwell. Cells were transfected as described for cGAS-STING reporter cell assays. To physically separate bacterial cells from reporter cells, transfected cGAS-STING reporter cells challenged by the addition of WT *H. pylori* 26695 added to either the apical transwell chamber (293T cells seeded directly into the cell culture well), or the basolateral compartment (293T cells seeded in the transwell apparatus) at a MOI of 20. As a control, cGAS-STING assays in which *H. pylori* were added directly to reporter cell monolayers were performed in parallel. After 24 h of bacterial infection, monolayers were processed as described for cGAS-STING activation assays, and luminescence values were obtained. IFN-β transcriptional reporter values were expressed as the normalized fold change over mock infected wells.

### STING transactivation assays

For STING transactivation assays, 293T cells were transfected with either (i) pUNO1-hcGAS-HA3X, pRL-SV40P, and pcDNA3.1 empty vector, or (ii) pUNO1-hSTING-R232, pRL-SV40P, and IFN-Beta-pGL3 using Lipofectamine 2000 as described for cGAS-STING activation assays. At 12 h post-transfection, cells were lifted using sterile phosphate buffered saline (PBS) and were re-plated in 24-well dishes at a 1:1 ratio at approximately 2 × 10^5^ cells/well. Cells were allowed to adhere and were subsequently challenged by the indicated *H. pylori* strain at a MOI of 20. After 24 h of bacterial infection, IFN-β transcriptional reporter activity was assayed as described for cGAS-STING activation experiments. STING transactivation by cGAMP diffusion was expressed as the normalized fold change of IFN-β transcriptional reporter activity over mock infected wells.

### TLR and RIG-I activation assays

To assess TLR9 or TLR7 activation, HEK293 stably transfected with hTLR9, hTLR7, or parental null1 cells (Invivogen) were seeded into 96-well plates at a density of approximately 2 × 10^4^ cells per well, and were subsequently by WT *H. pylori* 26695 or its isogenic mutant strains at a MOI of 100 in quadruplicate as previously described (Varga et al., 2016b; Varga et al., 2021). Supernatants were collected at 24 h post-infection, and TLR9 and TLR7 activation was determined by measuring secreted embryonic alkaline phosphatase (SEAP) reporter activity in cell culture supernatants by QuantiBlue^™^ reagent (Invivogen) using a microplate reader (BioTek Synergy HI) to record the absorbance at 650 nm. As a positive control for TLR7 activation, HEK293-hTLR7 reporter cells were stimulated with 5 μg/ml imiquimod (R837), an imidazoquinoline amine analog to guanosine (Invivogen). TLR9 or TLR7 activation was normalized to SEAP levels produced by infected null1 parental cells and is expressed as the fold-change over mock infected controls. For ciprofloxacin inhibition of bacterial DNA replication, ciprofloxacin was added to HEK293-hTLR9 co-cultures at the same time as bacterial inoculation at multiples of the previously reported minimum inhibitory concentration (MIC), with 1X MIC equivalent to 0.125 µg/ml. Inhibition of TLR9 activation is expressed as a percent of the normalized fold change over mock treated, *H. pylori*-challenged wells.

For RIG-I activation studies, HEK-Lucia™ hRIG-I cells were plated in a 96-well dish at approximately 5 × 10^4^ cells per well and were subsequently challenged by WT *H. pylori* 26695 or the indicated isogenic mutant strain at a MOI of 100 in quadruplicate. As a positive control, cells were transfected with 100 ng/ml of the RIG-I agonist 5’ triphosphate hairpin RNA (3p-hpRNA, Invivogen) complexed to Lipofectamine 2000 (Life Technologies) according to the manufacturer’s protocol. At 24 h post-challenge, RIG-I stimulation was assessed by analyzing Lucia luciferase reporter gene expression in 20 µl cell culture supernatants using QUANTI-Luc^™^ (Invivogen). Data are expressed as the normalized fold change over mock infected cells. TLR and RIG-I stimulation experiments were performed a minimum of three times with quadruplicate technical replicates per experimental condition.

### CagA translocation assays

AGS or 293T cells were plated in 12-well dishes at a seeding density of 1 × 10^5^ cells/well and were cultured overnight. Monolayers were challenged by the indicated *H. pylori* strain at a MOI of 100 for 6 h, as previously described (Shaffer et al., 2011; Varga et al., 2021). Wells were washed in sterile PBS to remove non-adherent bacteria, and cells were lysed in assay buffer (1% NP-40) supplemented with cOmplete™ mini EDTA-free protease inhibitor (Roche) and PhosSTOP phosphatase inhibitor (Roche). The soluble fraction was collected and prepared in 2X SDS buffer for immunoblotting. To assess CagA tyrosine phosphorylation, AGS or 293T samples were separated on 7.5% gels (Bio-Rad) for 60 min at 165V. Proteins were then transferred onto nitrocellulose using the TransBlot Turbo system (Bio-Rad) following manufacturer’s recommendations. Membranes were blocked in 3% BSA in Tris-buffered saline containing 0.1% Tween-20 (TBST), followed by incubation with anti-phosphotyrosine monoclonal antibody (*α*-PY99, Santa Cruz Biotechnology). Total CagA levels were assessed by subsequent incubation with an anti-CagA monoclonal antibody (*α*-CagA, Santa Cruz Biotechnology). Phosphorylation of CagA or total CagA was visualized using chemiluminescence (Pierce). TEM-CagA translocation was assayed as previously described (Varga et al., 2021).

### Bacterial adherence and internalization assays

Adherence and internalization into gastric or kidney epithelial cells were assessed as previously described(Varga et al., 2021). Briefly, WT *H. pylori* or the *cagX* isogenic derivative were co-cultured with AGS or 293T cells at a MOI of 100. After a 4 or 6 h infection, respectively, cell culture medium was aspirated, and cell monolayers were gently washed with sterile PBS to remove non-adherent bacteria. To assess intracellular bacteria, RPMI or DMEM supplemented with gentamycin (100 μg/mL) was added to a subset of wells, and the cells were incubated for an additional hour at 37°C in 5% CO_2_. To assess total adherent and intracellular bacteria, fresh RPMI or DMEM was replenished into the remainder of wells following the removal of non-adherent bacteria. After the 1 h incubation, RPMI or DMEM was aspirated and all wells were washed in sterile PBS, lysed by mechanical disruption, and were serially diluted on blood agar plates for colony enumeration. Experiments were performed a minimum of three times with triplicate technical replicates per cell line and culture condition.

### Detection of bacterial DNA in AGS cytosolic extracts

Digitonin extracts of AGS gastric epithelial cells were prepared essentially as previously described(West et al., 2015). Briefly, approximately 8.4 × 10^6^ AGS cells were infected with exponentially growing WT or *cagX H. pylori* at a MOI of 50. As a control, an equivalent number of AGS cells were mock infected. After 6 h of infection, cells were rinsed with sterile PBS and were treated with 50 units of Turbo DNaseI (Life Technologies) in digestion buffer at 37°C for 15 min. The cells were rinsed twice with PBS, trypsinized, and collected in 2 ml of sterile PBS. Collected cells were separated into aliquots of approximately 400 µl to generate ‘total’ cell extracts, and of approximately 1600 µl to generate ‘cytosolic’ extracts. Aliquots were centrifuged at 980 x g for 3 min and cell pellets were washed once with PBS. To generate the ‘total’ cell extract, one pellet for was re-suspended in 100 µl of 50 µM NaOH and incubated for 30 min at 95°C to solubilize DNA, followed by pH neutralization by the addition of 10 µl of 1 M Tris-HCl, pH 8. These extracts served as normalization controls for the quantitation of mitochondrial DNA (mtDNA) and bacterial DNA. To generate ‘cytosolic’ extracts, cell pellets were re-suspended in cytosolic extraction buffer (150 mM NaCl, 50 mM Tris pH 8.0, and 20 µg/ml Digitonin [Sigma-Aldrich]), and homogenates were rotated end-over-end for 10 min at room temperature for selective membrane permeabilization. Cytosolic fractions were separated from intact cells and nuclear/bacterial fractions by serial centrifugations at 17,000 x g for 3 min. Recovered supernatants were incubated for 10 min at 95°C to isolate DNA. To assess the presence of nuclear DNA, mtDNA, and *H. pylori* DNA in cellular fractions, standard PCR and qPCR targeting fragments of the *H. pylori* chromosome (*hp1421*, 290 bp), mtDNA (*coxII*, 444 bp(Fernandez-Moreno et al., 2016)), and nuclear DNA (*hNuc*, 467 bp (Fernandez-Moreno et al., 2016)) were amplified from ‘total’ and ‘cytosolic’ fractions using Taq polymerase (standard PCR) or Fast SYBR Green chemistry (qPCR) on a Viia7 platform (Thermo). *C*_T_ values obtained for cytosolic fractions were normalized to corresponding *C*_T_ values obtained for total cell extracts, and cytosolic enrichment of bacterial DNA was calculated as the normalized ratio of *hp1421 C*_T_ values to *coxII C*_T_ values. A minimum of four biological replicate experiments were performed for each experimental condition.

### Quantitation of secreted extracellular cGAMP

AGS cells seeded into T25 flasks were grown to approximately 80% confluency were mock infected or were challenged by *H. pylori* at a MOI of 100 for 6 h. Bacteria were inactivated by the addition of 200 µg/ml gentamicin, and monolayers were incubated overnight. At 24 h post-infection, equivalent volumes of cell culture supernatants were collected and concentrated via solvent evaporation. Samples were reconstituted in one-tenth volume assay buffer, and cGAMP levels were quantified using the DetectX® 2’,3’-cGAMP STING-Based FRET assay (Arbor Assays). For each experimental condition, cGAMP secretion assays were performed in triplicate and data represents a minimum of three biological replicates.

### Confocal laser scanning microscopy

Adult primary human gastric epithelial cells were grown on collagen-coated 12 mm glass coverslips (#1.5, 170 µm thickness) overnight prior to challenge by WT or *cagX H. pylori* at a MOI of 50. As a control, a subset of coverslips was mock infected. After 6 h, coverslips were washed in sterile PBS three times, followed by fixation in 4% paraformaldehyde in PBS for 20 min at room temperature. Coverslips were washed in PBS and cells were permeabilized in confocal blocking buffer (3% bovine serum albumin, 0.1% TritonX-100, 1% saponin in sterile PBS) for 1 h at room temperature. For immunostaining, coverslips were stained with anti-STING monoclonal antibody (Life Technologies, 1:100) in confocal blocking buffer overnight at 4°C. Coverslips were rigorously washed three times in PBST to remove unbound antibody and were subsequently incubated in AlexaFluor 488-conjugated secondary antibody (Life Technologies, 1:1000) in blocking buffer for 1 h at room temperature. For visualization of the nuclei and actin, samples were stained with stained with 4’,6-diamidino-2-phenylindole (DAPI) and AlexaFluor 594 phalloidin for 1 h at room temperature. Coverslips were washed in PBS and were mounted with ProLong Gold antifade (Life Technologies). Fluorescence images were captured using a 60X silicon immersion objective on an Olympus FV3000 confocal laser scanning microscope and images were acquired and processed using cellSens software (Olympus). Quantification of STING particle size and number was performed using Fiji software (ImageJ) with automated thresholding and subsequent particle analysis of segmented images for mock infected (7 fields of view, *n*=90 cells); WT infected (11 fields of view, *n*=82 cells); and *cagX* infected (9 fields of view, *n*=59 cells) gastric epithelial cells. To normalize across imaging conditions, average particle sizes were calculated by multiplying the average pixel area by the pixel resolution for each field of view.

### RNA isolation and gene expression analyses

Primary gastric epithelial cells were mock infected (*n*=3 biological replicates) or co-cultured with WT *H. pylori* 26695 (*n*=6 biological replicates) or the corresponding *cagX* isogenic mutant (*n*=3 biological replicates) for 6 h. Cells were washed three times with sterile PBS and total RNA was isolated using the Direct-Zol Miniprep Plus kit (Zymo Research) following the manufacturer’s protocol. Total RNA was stringently digested with Turbo DNase I (Invitrogen) to remove contaminant DNA. To determine inflammatory gene expression changes in response to *H. pylori* infection, DNA-free RNA (100 ng per sample) was analyzed on the nCounter Sprint Profiler (NanoString Technologies) using the nCounter Human Host Response Panel (NanoString Technologies), which simultaneously quantifies transcripts for 773 immune-related genes and 12 internal reference genes. Differences in gene expression between experimental groups was calculated using the ROSALIND Platform for nCounter Analysis (https://rosalind.onramp.bio/). Raw data (RCC files) were normalized to internal reference genes the nSolver 4.0 software integrated within the ROSALIND platform (ROSALIND, Inc.). Gene normalization was performed using housekeeping probes selected based on the geNorm algorithm as implemented in the Bioconductor package NormqPCR. Differentially expressed genes in *H. pylori* challenged cells were determined using the Benjamini– Hochberg *P* value adjustment method of estimating false discovery rates (FDR), with significance set at *p*<0.05. Read distribution percentages, violin plots, identity heatmaps, and sample multidimensional scaling (MDS) plots were generated within ROSALIND during sample QC. Read normalization, gene expression fold changes, and the associated *P* values were calculated using criteria provided by NanoString. Pathway enrichment analysis was performed within ROSALIND using the REACTOME database, and gene term enrichment was calculated using a hypergeometric distribution algorithm in reference to the background set of genes in the panel with significance set at *p*<0.01 and greater than ± 1.8-fold gene expression enrichment. Volcano plots and heat maps were generated in GraphPad Prism using normalized gene expression data exported from ROSALIND. Volcano plots were constructed by plotting the log_2_ of the normalized fold change versus the −log_10_ of the adjusted *P* value for each gene. The dashed *P*_adj_ lines demarcate genes meeting the threshold for significance (*P*_adj_<0.01 and *P*_adj_<0.05) after correction with the Benjamini–Hochberg procedure for controlling FDR.

### Gastric epithelial cell supernatant transfer assay

Primary gastric epithelial cells seeded into 24-well gelatin-coated dishes were mock infected or were co-cultured with WT *H. pylori* or the *cagX* isogenic mutant at a MOI of 100 for 6 h or 12 h. Cell culture supernatants were harvested and treated with 100 µg/ml gentamicin for 1 h at 37°C to eliminate viable bacteria. Supernatants were subsequently spun for 10 min at 10,000 x g to remove bacteria and gastric cells, and cleared supernatants were stored at −20°C. For TLR9 stimulation studies, HEK293-hTLR9 cells were seeded into 96-well plates at a density of approximately 2 × 10^4^ cells per well in a volume of 100 µl DMEM per well. Primary gastric epithelial cell supernatants were added to hTLR9 reporter cells at an equal volume and incubated for 24 h at 37°C in 5% CO_2_. TLR9 activation was determined via SEAP levels determined by QuantiBlue^™^ (Invivogen) using a microplate reader (BioTek Synergy HI) to record the absorbance at 650 nm. The concentration of cell-free DNA in processed supernatants was determined by high sensitivity dsDNA Qubit assay (Life Technologies).

For enzyme treatments of gastric epithelial cell supernatants, pre-cleared supernatants obtained from 6 h *H. pylori* challenged gastric epithelial cells were either treated by (i) the addition of 10 units Turbo DNase (Life Technologies) and incubation at 37°C for 30 min, (ii) heating the supernatant to 70°C for 15 min, or (iii) heating the supernatant to 70°C for 15 min followed by cooling to room temperature and subsequent DNase treatment as described in (i). Treated supernatants were added to HEK293-hTLR9 reporter cells as described for supernatant transfer assays, and TLR9 stimulation was assessed by QuantiBlue^™^ after 24 h incubation. TLR9 activation is expressed as the fold change over mock treated HEK293-hTLR9 cells.

### Extracellular vesicle (EV) purification

Primary gastric epithelial cells were challenged by *H. pylori* or were mock infected for 6 h. Supernatants were pre-cleared by serial centrifugations (10,000 x g) at 4°C. EV-containing supernatants (2 ml per biological replicate) were subsequently magnetically labeled for 1 h at room temperature by CD9, CD63, and CD81 antibodies (Human Exosome Isolation Kit, Pan, Miltenyi Biotec) followed by EV isolation via magnetic separation and elution. Immunoaffinity purified exosomes were subjected to qPCR analysis probing for *H. pylori* genomic DNA (*hp1421* locus, 290 bp fragment) and mtDNA (*coxII*, 444 bp fragment) by Fast SYBR Green chemistry (Life Technologies) on a Viia7 platform (Thermo). *H. pylori* DNA enrichment within purified EVs was determined by quantifying the ratio of *hp1421 C*_T_ values to *coxII C*_T_ values for EVs obtained from WT and *cagX*-infected gastric cells. EVs purified from mock infected gastric epithelia contained levels of mtDNA similar to EVs derived from *H. pylori*-challenged primary cells.

### Immunoprecipitation of cGAS-DNA complexes

293T cells were grown in 6-well plates for 24-30 hours to achieve approximately 90% confluency. Cells were transfected with pUNO1-hcGAS-HA3X (240 ng DNA/well) complexed to Lipofectamine 2000. After 12 – 16 h, approximately 7.2 × 10^6^ transfected 293T cells were challenged by exponentially growing WT or *cagX H. pylori* at a MOI of 100. As a control, an equivalent number of transfected 293T cells were mock infected. After 6 h of infection, 293T cells were rinsed with PBS and DNA-protein complexes were cross-linked by 1% paraformaldehyde for 15 min at room temperature. Cross-linking reactions were quenched by the addition of 250 mM glycine, and cells were collected by mechanical detachment and centrifugation at 4000 rpm for 10 min. Cell pellets were washed once with sterile PBS, followed by re-suspension in pre-chilled lysis buffer (5 mM EDTA, 1% NP-40, and 1X protease inhibitor cocktail in PBS) and sonication at 5% amplitude (10 Sec ON, 10 Sec OFF, 4-6 cycles) to generate cleared lysates. Sonicated cell extracts were centrifuged at 14,000 rpm for 30 min at 4°C. The recovered cell-free supernatant was incubated with 4 µg of anti-HA monoclonal antibodies (clone 12CA5) overnight at 4°C with continuous end-over-end rotation. The following day, 50 µl of Protein G Dynabeads (Life Technologies) pre-blocked in PBS containing 1% BSA were incubated with immunopurification samples for 90-120 min at 4°C with continuous end-over-end rotation. Dynabeads were collected by magnetic isolation, washed twice with 1X cell lysis buffer, followed by one wash in high salt wash buffer (cell lysis buffer + 300 mM NaCl). Magnetic beads were re-suspended in 100 µl of 1% SDS + 0.1 M sodium bicarbonate buffer and de-crosslinked by incubation at 60°C overnight. Purification of cGAS-HA3x was confirmed in the eluted fractions by immunoblot analysis. To purify DNA complexed with cGAS-HA3x, de-crosslinked fractions were treated with 20 µg of Proteinase K (Sigma-Aldrich) at 60°C for 1-2 h, followed by DNA isolation via the ChIP DNA Clean and concentrator kit (Zymo Research) according to the recommended protocol. Eluted DNA was quantified using the Qubit high sensitivity dsDNA assay (Life Technologies).

### cGAS ‘ChIP-seq’ library preparation and sequencing

ChIP samples were quantified using Qubit 2.0 Fluorometer (Life Technologies) the DNA integrity was analyzed with 4200 TapeStation (Agilent Technologies). cGAS ‘ChIP-seq’ library preparation and sequencing reactions were conducted at GENEWIZ, Inc. (South Plainfield, NJ, USA). NEB NextUltra DNA Library Preparation kit was used following the manufacturer’s recommendations (Illumina). Briefly, DNA eluted from cGAS immunopurifications was end repaired and adapters were ligated after adenylation of the 3’ ends. Adapter-ligated DNA was size selected, followed by clean up, and limited cycle PCR enrichment. The cGAS ‘ChIP’ library was validated using Agilent TapeStation and quantified using Qubit 2.0 Fluorometer as well as RT-PCR (Applied Biosystems). Sequencing libraries were multiplexed and clustered on two lanes of a flowcell. After clustering, the flowcell was loaded on the Illumina HiSeq instrument according to manufacturer’s protocol (Illumina). Sequencing was performed using a 2×150 paired end (PE) configuration. Image analysis and base calling were conducted by the HiSeq Control Software (HCS). Raw sequence data generated from Illumina HiSeq was converted into fastq files and de-multiplexed using Illumina’s bcl2fastq 2.17 software. One mismatch was allowed for index sequence identification. Sequence reads were processed to remove adapter sequences and nucleotides with poor quality at both 5’ and 3’ ends using CLC Genomics workbench. Sequence reads below 15 bases were discarded. Trimmed data was aligned to both human (*Homo sapiens* reference genome hg38) and *H. pylori* 26695 (reference genome NC_000915) reference genomes. Only specific alignment was allowed during mapping. To detect peaks that were differentially present in cGAS purifications obtained from WT-infected cells versus *cagX*-infected cells, reads were normalized to mock infected control preparations, and peak calling was performed using the Transcription Factor model within CLC Genomics workbench with *p*<0.01 considered significant.

### Nanoluciferase (NanoLuc) bioluminescence assays

AGS or 293T cells were cultured overnight to reach approximately 70% confluence, and were challenged by WT or *cagX H. pylori* [NanoLuc] (26695) or WT or *cagE H. pylori* [NanoLuc] (G27) at a MOI of 50 for 24 h. Cell culture supernatants were removed and monolayers were washed in sterile PBS to remove non-adherent cells. Monolayers were lysed in 1% NP-40, and 50 µl of cell lysate was transferred to a white walled 96-well plate. To measure nanoluciferase bioluminescence, 20 µl Nano-Glo Luciferase substrate assay buffer containing furimazine (Promega) prepared according to the manufacturer protocol was added to each well. Luciferase activity was immediately assessed using a BioTek Synergy H1 plate reader with luminesce acquisition settings set as recommended by the manufacturer, with the exception of the gain which was adjusted to 230. To determine the level of background NanoLuc activity produced by *H. pylori*, 20 µl of overnight bacterial cultures were directly lysed in Nano-Glo Luciferase substrate assay buffer containing furimazine, and luciferase values were immediately obtained via plate reader using the same parameters as for eukaryotic cells. Background luminescence produced by *H. pylori* was determined by normalizing luciferase values by the corresponding culture OD_600_ and is expressed as the fold change in luminescence over values obtained for *H. pylori* cultures that do not harbor the nanoluciferase expression construct. Bioluminescence of *H. pylori*-challenged wells is expressed as the fold change over mock infected wells. For each eukaryotic cell line, a minimum of four biological replicate experiments were performed in 24-well plate technical replicate format.

### Live cell fluorescence microscopy analysis of LifeAct-mScarlet

*H. pylori* harboring LifeAct-mScarlet constructs were grown to exponential phase in broth culture and AGS or 293T monolayers were inoculated at a MOI of 50 overnight at 37°C in 5% CO_2_. Monolayers were washed in sterile PBS to remove non-adherent bacteria, and monolayers were imaged via live cell, phase contrast epi-fluorescence microscopy on a Nikon Ti Eclipse equipped with a 594_em_ filter and a Plan Apo VC 20X/0.75 NA air objective. Fluorescent images were superimposed on the corresponding phase contrast image of the same field of view. Images were processed for equivalent contrast, brightness, and magnification using the OMERO platform (Allan et al., 2012).

### Flow cytometry analysis

AGS or 293T cells were cultured overnight to reach approximately 70% confluence, and were challenged by WT or *cagX H. pylori* [LifeAct-mScarlet] at a MOI of 50 for 24 h. Cell culture supernatants were removed and monolayers were washed in sterile PBS to remove non-adherent cells. Cells were trypsinized (AGS) or mechanically detached (293T) from tissue culture flasks, washed once in PBS via centrifugation and pelleting, and re-suspended in PBS at approximately 1 × 10^6^ cells/ml. Samples were analyzed on an Attune^™^ NxT Flow Cytometer (Thermo). Forward scatter-height (FSC-H) and sideward scatter-height (SSC-H) profiles were used in gating strategies to select for single cells, and positive mScarlet fluorescence gates were determined by analyzing 293T cells that had been transfected with pLifeAct_mScarlet-i_N1 (Bindels et al., 2017).

### IL-8 quantitation

Interleukin-8 (IL-8) secretion was monitored by human CXCL8 ELISA (R&D Systems) as previously described(Shaffer et al., 2011; Varga et al., 2021). Briefly, AGS cells were plated in 24-well dishes and were cultured overnight prior to infection with *H. pylori* at a MOI of 100 for 4.5 h. Supernatants were collected and stored at −20°C until analysis by ELISA. A minimum of three biological replicate experiments were performed in triplicate for all strains, and IL-8 secretion is expressed as a percent of IL-8 levels induced by WT *H. pylori* for each replicate experiment.

### ‘Transfer DNA’ immunoprecipitation

To assess whether processed chromosomal DNA fragments physically associate with the *cag* T4SS outer membrane complex, *H. pylori* grown for 24 h on blood agar were harvested in 2 ml PBS, and 50 µL of the cell suspension were removed to serve as the ‘input’. The remainder of the collected cells were pelleted by centrifugation and were washed once in PBS. Cell pellets were re-suspended in 500 µL PBS, and protein-DNA complexes were cross-linked by the addition of 500 µL 1% paraformaldehyde for 10 min at room temperature, followed by quenching with 1 ml 250 mM glycine. Cross-linked cells were pelleted and washed once with PBS. Cell pellets were re-suspended in 1 mL lysis buffer (5 mM EDTA, 1% NP-40 in PBS) supplemented with 2X cOmplete™ inhibitor (Roche) and were sonicated until the lysate became turbid. Cell lysates were treated with 2 units of Turbo DNase (Life Technologies) in 10 mM MgCl_2_ for 30 min at room temperature. To solubilize membranes, 0.2% SDS and 0.2% sodium deoxycholic acid (final concentrations) were added to cell lysates, and samples were rotated end-over-end for 1-2 h. Supernatants were separated from insoluble cell debris by centrifugation at 14,000 rpm for 30 minutes at 4°C. In a separate tube, 4 µl polyclonal anti-CagY antisera (a kind gift from Dr. Tim Cover) was added to 800 µl lysis buffer containing 10 mM MgCl_2_, 2 units Turbo DNase, and 25 µl Protein G Dynabeads, and was incubated for 10 min at room temperature. Cleared supernatants were added directly to CagY antibody solutions and were incubated for an additional 2-4 h with continuous end-over-end rotation. Beads were isolated by magnetic separation and were washed twice in PBS supplemented with 10 mM MgCl_2_ and 1 unit Turbo DNase, followed by two washes in high salt buffer (lysis buffer containing 400 mM NaCl), and a final wash in PBS. Protein-DNA complexes were eluted and de-cross-linked in 100 µl 1% SDS in 0.1 M NaHCO_3_ at 65°C overnight. The following day, proteins were digested using Proteinase K (10 µg) at 65°C for 30 minutes. DNA was precipitated by 100% ethanol in 0.3 M sodium acetate (1:3 v/v) at −20°C. Precipitated DNA was pelleted by centrifugation at 14,000 rpm for 30 minutes at 4°C, washed once with 70% ethanol, and re-hydrated in 20 µl ultrapure water. To serve as a control, the initial ‘input’ cell pellet was re-suspended in 100 µl of 50 µM NaOH, and DNA was liberated by incubating at 95°C 30 min, followed by pH neutralization by the addition of 10 µl of 1M Tris, pH 8. To assess chromosomal DNA association with immunopurified CagY complexes, 1 µl of ‘input’ and ‘IP’ DNA samples were used as the template in standard PCR assays targeting a 795 bp chromosomal DNA amplicon. Quantitation of DNA amplification from ‘input’ and ‘IP’ samples was conducted by densitometry analysis and amplification efficiency of ‘IP’ samples was calculated as a percent of the corresponding ‘input’ sample amplification for each biological replicate experiment. Transfer DNA immunopurification assays were performed a minimum of four times per strain.

### Statistical analyses

Data are expressed as mean values ± standard error of the mean, which were calculated from a minimum of three biological replicate experiments. In all graphs, each data point represents an individual measurement, lines represent the mean, and error bars represent the standard error of the mean. Statistical analyses were performed using GraphPad Prism 9 software, with differences of *p*<0.05 considered statistically significant.

